# Bayesian multivariate reanalysis of large genetic studies identifies many new associations

**DOI:** 10.1101/638882

**Authors:** Michael C. Turchin, Matthew Stephens

## Abstract

Genome-wide association studies (GWAS) have now been conducted for hundreds of phenotypes of relevance to human health. Many such GWAS involve multiple closely-related phenotypes collected on the same samples. However, the vast majority of these GWAS have been analyzed using simple univariate analyses, which consider one phenotype at a time. This is despite the fact that, at least in simulation experiments, multivariate analyses have been shown to be more powerful at detecting associations. Here, we conduct multivariate association analyses on 13 different publicly-available GWAS datasets that involve multiple closely-related phenotypes. These data include large studies of anthropometric traits (GIANT), plasma lipid traits (GlobalLipids), and red blood cell traits (HaemgenRBC). Our analyses identify many new associations (433 in total across the 13 studies), many of which replicate when follow-up samples are available. Overall, our results demonstrate that multivariate analyses can help make more effective use of data from both existing and future GWAS.

**Author Summary:** Genome-wide association studies (GWAS) have become a common and powerful tool for identifying significant correlations between markers of genetic variation and physical traits of interest. Often these studies are conducted by comparing genetic variation against single traits one at a time (‘univariate’); however, it has previously been shown that it is possible to increase your power to detect significant associations by comparing genetic variation against multiple traits simultaneously (‘multivariate’). Despite this apparent increase in power though, researchers still rarely conduct multivariate GWAS, even when studies have multiple traits readily available. Here, we reanalyze 13 previously published GWAS using a multivariate method and find >400 additional associations. Our method makes use of univariate GWAS summary statistics and is available as a software package, thus making it accessible to other researchers interested in conducting the same analyses. We also show, using studies that have multiple releases, that our new associations have high rates of replication. Overall, we argue multivariate approaches in GWAS should no longer be overlooked and how, often, there is low-hanging fruit in the form of new associations by running these methods on data already collected.

## 2 Introduction

Genome wide association studies (GWAS) have been widely used to identify genetic factors – particularly single nucleotide polymorphisms (SNPs) and copy number variations (CNVs) – associated with human disease risk and other phenotypes of interest (Price et al., 2015; Visscher et al., 2017). Indeed, at time of writing over 24,000 such associations have been identified as ‘genome-wide significant’ (MacArthur et al., 2017).

The vast majority of these many genetic association analyses consider only one phenotype at a time (“univariate association analysis”). This is despite the fact that measurements on multiple phenotypes are often available, and joint association analysis of multiple phenotypes (“multivariate association analysis”) can substantially increase power (Jiang and Zeng, 1995; Zhu and Zhang, 2009; Shriner, 2012; Yang and Wang, 2012; Galesloot et al., 2014). There are likely multiple reasons for the preponderance of univariate analyses. One possible reason is that initial association analyses are usually performed under tight time constraints, and at a time when many other analysis issues (e.g. quality control, population stratification) are competing for attention. In these conditions it makes sense to focus on the simplest possible approach that will quickly yield new associations, without overly worrying about loss of efficiency. In addition analysts may be legitimately concerned that deviation from the most widely adopted analysis pipeline may invite unwanted additional reviewer attention.

Nonetheless, we believe that multivariate association analysis has an important role to play in making the most of costly and time-consuming GWAS studies. One way forward is to conduct multivariate analyses of previously-published GWAS, checking for additional associations that may have been missed by the initial univariate association analyses. This is greatly facilitated by the fact that many GWAS now make summary data from single-SNP tests freely available (Willer et al., 2013; Wood et al., 2014; Locke et al., 2015; Shungin et al., 2015; Astle et al., 2016), and that simple multivariate analysis can be conducted using such summary data (Stephens, 2013; Pickrell et al., 2016; Hormozdiari et al., 2016).

Here we demonstrate the potential benefits of reanalyzing published GWAS using multivariate methods. Specifically we apply multivariate methods from Stephens 2013 to reanalyze 13 different GWAS whose initial publications reported only univariate results. In most cases our multivariate analyses find many new associations. For example, in GIANT 2014/5 we find over 150 new associations. In studies with multiple data releases, we find that new multivariate associations found in initial releases typically replicate in subsequent releases, supporting that many of the new associations are likely real. We also demonstrate that the multivariate approach is not equivalent to simply relaxing the univariate GWAS significance threshold. Finally, we point out some limitations of the specific framework we used here, and suggest some alternative strategies that may help address those limitations in future multivariate GWAS analyses.

## 3 Results

### Multivariate association analyses

To facilitate multivariate association analyses using the methods from Stephens 2013, we implemented them in an R package bmass (Bayesian multivariate analysis of summary statistics). The software requires as input univariate GWAS summary statistics, for the same set of SNPs, on *d* related phenotypes. Then, for each SNP, it attempts to categorize each phenotype as belonging to one of three categories: **U**nassociated, **D**irectly Associated, or **I**ndirectly Associated with the SNP. The difference between **D** and **I** is that an indirect association disappears after controlling for associations with other phenotypes (see Online Methods and Supplementary Figure 1).

For *d* phenotypes, there are 3*^d^* possible assignments of phenotypes to these 3 categories, and each assignment corresponds to a different “model” *γ*. For example, one model corresponds to the “null” that all phenotypes are **U**nassociated; another model corresponds to the model that all phenotypes are **D**irectly associated; another model corresponds to just the first phenotype being **D**irectly associated, etc. The goal of the association analysis is to determine which of these models is consistent with the data and, in particular, to assess overall evidence against the null model.

The support in the data for model *γ*, relative to the null model, is summarized by a Bayes Factor (BF*_γ_*). Large values of BF*_γ_* indicate strong evidence for model *γ* compared against the null. One advantage of Bayes Factors over *p*-values is that the Bayes Factors from different models can be easily compared and combined. For example, the overall evidence against the null is given by the (weighted) average of these BFs:

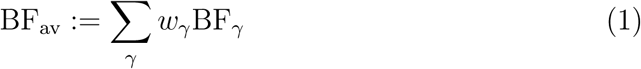

where the weights *w_γ_* are chosen to reflect the relative plausibility of each model *γ*. In bmass we implemented the Empirical Bayes approach from Stephens 2013 that learns appropriate weights from the data (see Online Methods).

### Comparisons with published univariate analyses

To provide a benchmark against which to compare our multivariate analysis results, we compiled a list of “previous univariate associations”: SNPs that were both reported as significant in the original publication and exceeded the original publication’s definition for genome-wide significance in at least one phenotype in the publicly-available (univariate) summary data analyzed here. This does not include all SNPs reported in every original publication because in some studies SNPs became genome-wide significant only after adding additional samples not included in the publicly available summary data.

We used these previous univariate associations to determine a significance threshold for our multivariate associations. Specifically, we declared a multivariate association as significant if its BF_av_ exceeds that of any previous univariate association’s BF_av_ in the same study (Stephens, 2013). The rationale is that the evidence for these multivariate associations exceeds the evidence for previously-reported genome-wide significant associations, which are generally regarded as likely to be (mostly) real associations.

Finally, we defined a list of “new multivariate associations”, which are SNPs that are significant in our multivariate analysis but are not a “previous univariate association”. To avoid double-counting of signals due to linkage disequilibrium (LD), we pruned the list of new multivariate associations so that they are all at least 0.5Mb apart. For additional details, see Online Methods.

### Many new loci identified in reanalyzing 13 publicly available GWAS studies

We applied bmass to 13 publicly available GWAS studies, representing 10 different collections of phenotypes (Table 1). Phenotypic collections include blood lipid traits (GlobalLipids: (Teslovich et al., 2010; Willer et al., 2013)), body morphological traits (GIANT: (Lango Allen et al., 2010; Speliotes et al., 2010; Heid et al., 2010; Wood et al., 2014; Locke et al., 2015; Shungin et al., 2015)), red blood cell traits (HaemgenRBC: (van der Harst et al., 2012; Astle et al., 2016)), blood pressure traits (International Consortium for Blood Pressure Genome-Wide Association et al., 2011; Wain et al., 2011), bone density traits (Zheng et al., 2015), and kidney function traits (Kottgen et al., 2010; Boger et al., 2011). For three of these phenotypic collections (GlobalLipids, GIANT, and HaemgenRBC), two different releases were available from the source consortiums. We conducted basic QC as described in Online Methods.

**Table 1:**
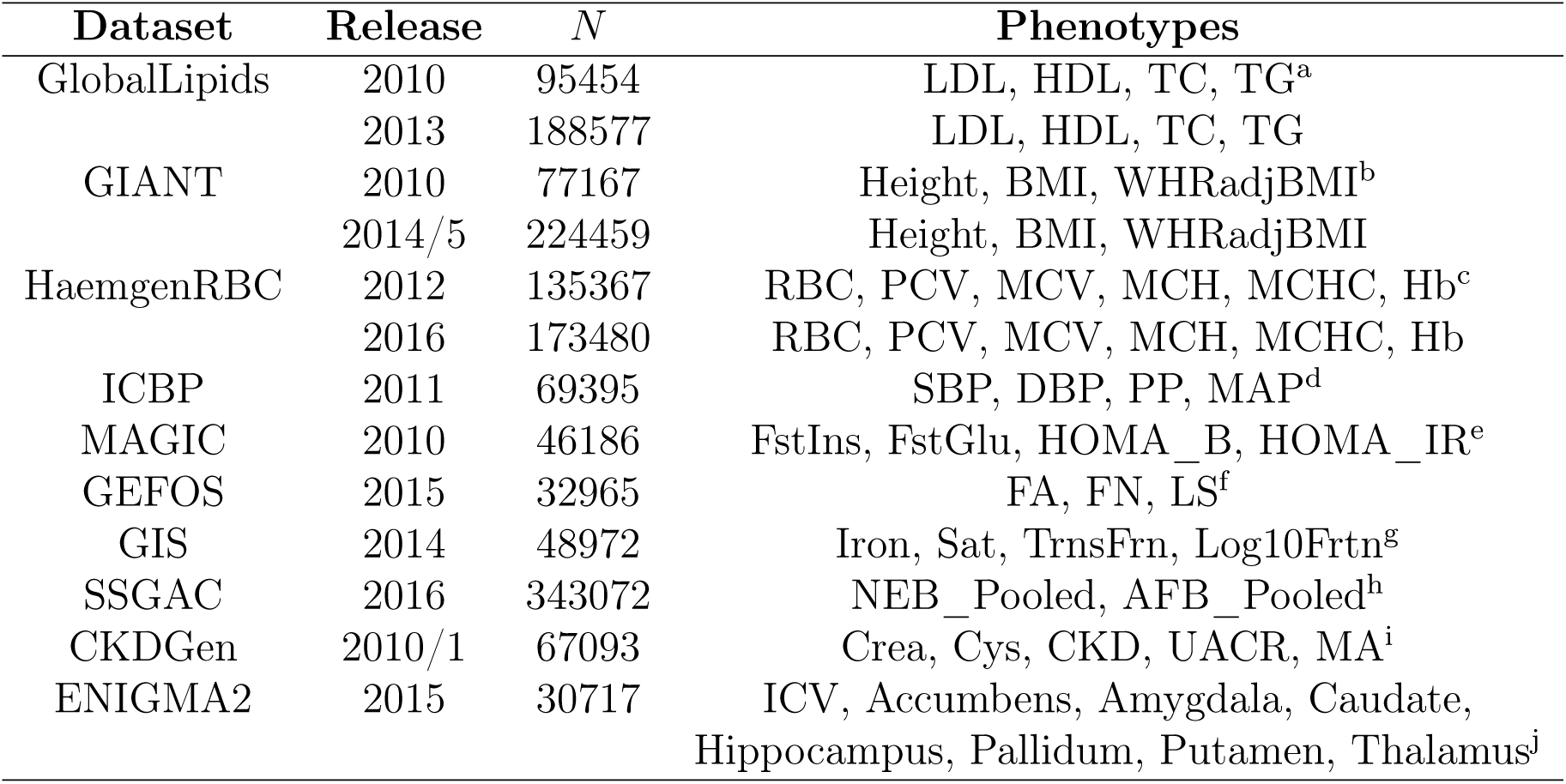
Dataset Summary. N is the maximum number of samples contributing to each study. a - Low-Density Lipoproteins (LDL), High-Density Lipoproteins (HDL), Total Cholesterol (TC), Total Triglycerides (TG) b - Body Mass Index (BMI), Waist-Hip Ratio adjusted for BMI (WHRadjBMI) c - Red Blood Cell Count (RBC), Packed Cell Volume (PCV), Mean Cell Volume (MCV), Mean Cell Haemoglobin (MCH), Mean Cell Haemoglobin Concentration (MCHC), Haemoglobin (Hb) d - Systolic Blood Pressure (SBP), Diastolic Blood Pressure (DBP), Pulse Pressure (PP), Mean Arterial Pressure (MAP) e - Fasting Insulin (FstIns), Fasting Glucose (FstGlu), Homeostatic Model Assessment of Beta Cell Function (HOMA_B), Homeostatic Model Assessment of Insulin Resistance Function (HOMA_IR) f - Forearm Bone Mineral Density (FA), Femoral Neck Bone Mineral Density (FN), Lumbar Spine Bone Mineral Density (LS) g - Serum Iron (Iron), Serum Transferrin Saturation (Sat), Serum Transferrin (TrnsFrn), Log-Transformed Ferritin (Log10Frtn) h - Number of Children Ever Born, Male & Female (NEB_Pooled), Age at First Birth, Male & Female (AFB_Pooled) i - Serum Creatine (Crea), Serum Cystatin (Cys), Chronic Kidney Disease (CKD), Urinary Albumin-to-Creatine Ratio (UACR), Microalbuminuria (MA) j - Intracranial Volume (ICV), specified subcortical brain structures refer to MRI-derived volume measurements for each one

| Dataset | Release | *N* | Phenotypes |
| --- | --- | --- | --- |
| GlobalLipids | 2010 | 95454 | LDL, HDL, TC, TG<sup>a</sup> |
|  | 2013 | 188577 | LDL, HDL, TC, TG |
| GIANT | 2010 | 77167 | Height, BMI, WHRadjBMI<sup>b</sup> |
|  | 2014/5 | 224459 | Height, BMI, WHRadjBMI |
| HaemgenRBC | 2012 | 135367 | RBC, PCV, MCV, MCH, MCHC, Hb<sup>c</sup> |
|  | 2016 | 173480 | RBC, PCV, MCV, MCH, MCHC, Hb |
| ICBP | 2011 | 69395 | SBP, DBP, PP, MAP<sup>d</sup> |
| MAGIC | 2010 | 46186 | FstIns, FstGlu, HOMA_B, HOMA_IR<sup>e</sup> |
| GEFOS | 2015 | 32965 | FA, FN, LS<sup>f</sup> |
| GIS | 2014 | 48972 | Iron, Sat, TrnsFrn, Log10Frtn<sup>g</sup> |
| SSGAC | 2016 | 343072 | NEB_Pooled, AFB_Pooled<sup>h</sup> |
| CKDGen | 2010/1 | 67093 | Crea, Cys, CKD, UACR, MA<sup>i</sup> |
| ENIGMA2 | 2015 | 30717 | ICV, Accumbens, Amygdala, Caudate, Hippocampus, Pallidum, Putamen, Thalamus<sup>j</sup> |

Our multivariate analyses identify, in total, hundreds of new associations. The numbers of previous univariate associations and new multivariate associations are summarized in Figure 1 (see also Supplementary Table 2). For example, we identify 162 new multivariate associations in GIANT2014/5, 65 in GlobalLipids2013, and 60 in HaemgenRBC2016. These represent power increases from 10% to 45% compared with previous univariate analyses.

**Figure 1:**
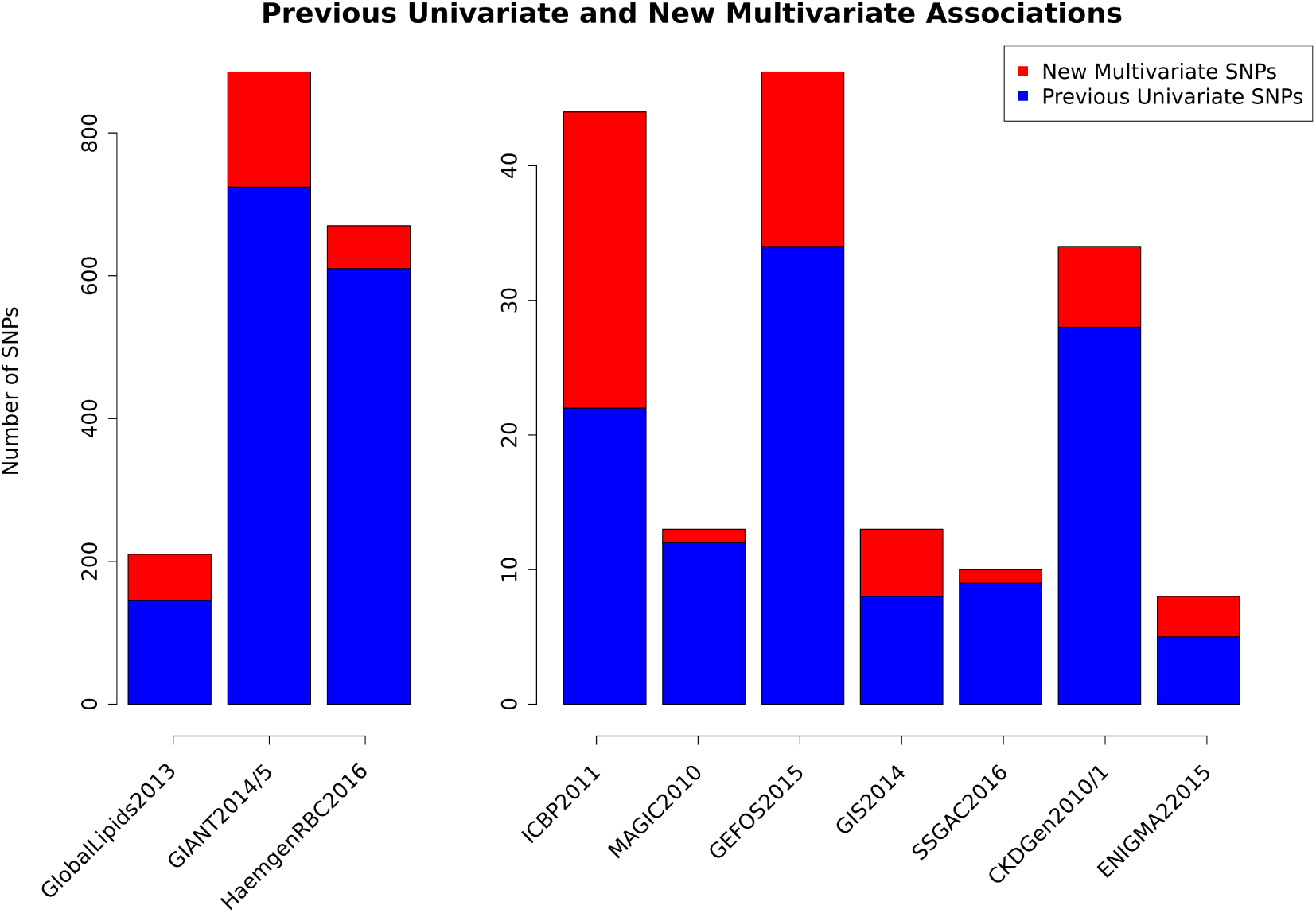
Number of Independent Significant SNPs, By Study. The barplot shows the number of independent SNPs that were significant in previous univariate analyses (blue) and the number of additional significant associations in our new multivariate analyses (red). For univariate analysis, significance levels were set by the original study. For multivariate analyses, we declared a SNP to be significant if its weighted average Bayes Factor (BF_av_) exceeded that of the smallest BF_av_ among the previous univariate significant SNPs. We considered SNPs more than .5Mb apart to be independent. See Table 1 and Online Methods for phenotype details, Online Methods for further analysis details, and Supplementary Tables 2-4 for lists of significant SNPs from each dataset.

### Replication of multivariate associations across releases

To demonstrate that many of these new multivariate associations are likely to be real we take advantage of three datasets that each have two releases separated by several years (GlobalLipids, GIANT, and HaemgenRBC). In each case we performed multivariate association analysis of the earlier release and checked how the new multivariate associations fared in univariate analyses of the later release (Figure 2). Since later releases include the samples from earlier releases, to assess “replication” we focus on whether the association in the new release is more significant than the original release – that is, whether the signal in the new (non-overlapping) samples provides additional evidence *over and above* the original signal. By this measure the results show high replication rates for the new multivariate associations: in total, 84 of 94 new associations have smaller minimum univariate *p*-values in the later release (at exactly the same SNP), and indeed the majority of these reach univariate GWAS significance in the later release.

**Figure 2:**
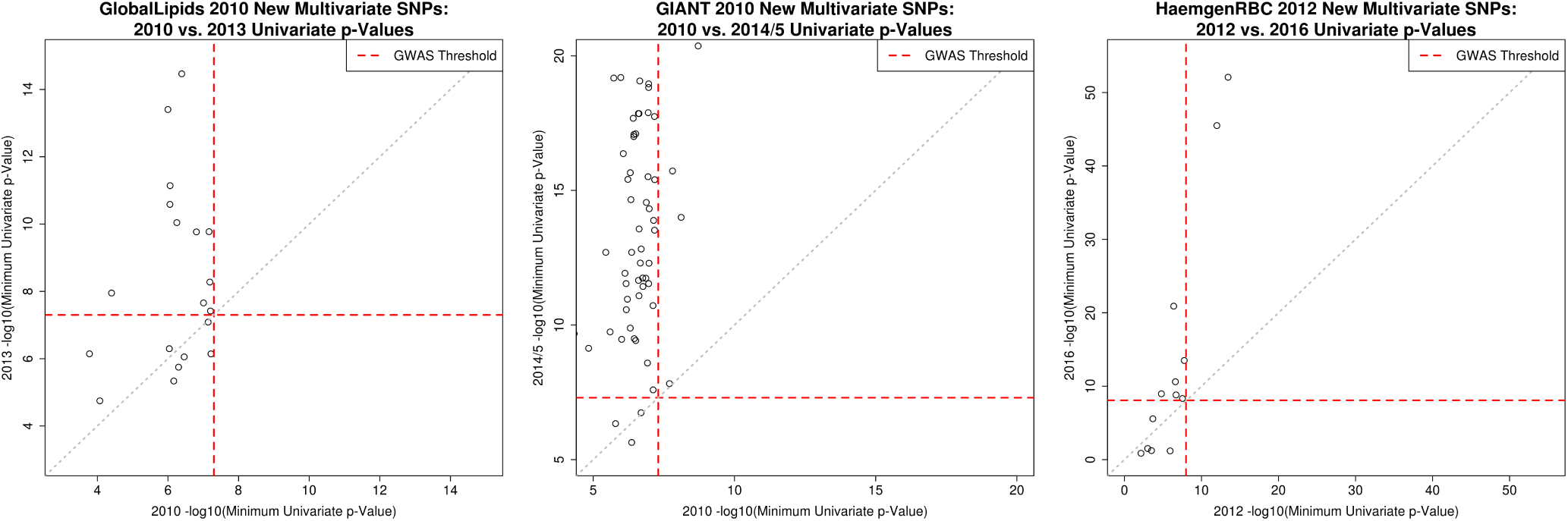
Replication of New Multivariate Associations. The figure shows results based on earlier and later releases from studies with multiple releases (GlobalLipids, GIANT, and HaemgenRBC). Each point represents a new multivariate association identified in our multivariate analysis of the earlier release. The *x*- and *y*-axes show the minimum (across phenotypes) of the -log_10_ univariate *p*-values from the earlier release (*x*-axis) vs. the later release (*y*-axis). Dashed red lines represent the univariate significance GWAS thresholds used for each study’s releases. Across all three studies, 84 out of 94 new multivariate associations from the earlier releases have smaller minimum univariate *p*-values in the later release, and 68 out of 84 new multivariate associations that did not reach GWAS significance in the earlier release do so in the later release (see Supplementary Table 5 for a per-dataset breakdown).

### Multivariate analysis is different from multiple univariate analyses

Because multivariate analysis takes account of *joint* patterns across phenotypes, its ranking of significance of SNPs can change compared with that from the univariate *p*-values alone. That is, multivariate analysis is not simply equivalent to multiple univariate analyses. To illustrate this we examined, in three well-powered studies, the associations that would be declared significant if the univariate significance threshold were relaxed, and assessed which of them would also be significant in our multivariate analysis (i.e. we assess whether, if we go deeper into the univariate results, we find the same SNPs as appear in our multivariate results). The results are shown in Figure 3. Although there is, understandably, substantial overlap between the significant SNPs, any non-trivial relaxation of the univariate threshold includes many SNPs that are not multivariate significant in our analysis; for example, at a univariate threshold of 5*×* 10*^−^*^7^ only 66% of the univariate significant SNPs are also multivariate significant across these three studies. This demonstrates that, indeed, our multivariate approach reorders significance of SNPs compared with multiple univariate analyses.

**Figure 3:**
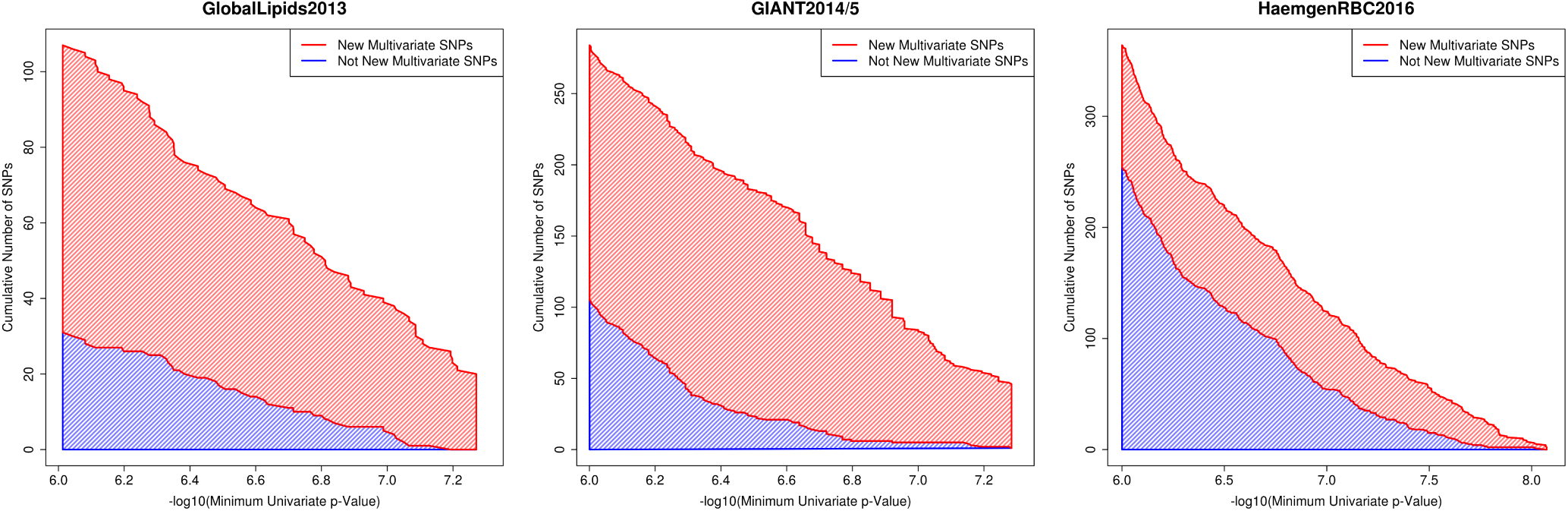
Comparison of New Multivariate Hits vs. Relaxing Univariate *p*-Value Threshold. For each data set the graph shows how many associations become significant as the univariate *p*-value threshold is relaxed (moving from right to left on the *x*-axis), and how many of these are declared as new multivariate hits in our analysis. In both cases results are pruned to avoid counting associations of SNPs in strong LD; see Online Methods for details. The appearance of appreciable blue areas indicates that the multivariate analysis is reordering the significance of SNPs compared with performing multiple univariate analyses.

### Reanalysis also identifies new univariate associations

During our multivariate reanalyses we noticed many SNPs that appeared to be genome-wide univariate significant but were – somewhat mysteriously – not reported as such by the original studies (i.e. SNPs whose univariate *p*-values crossed the significance threshold, as defined by the given study, in at least one trait). Supplementary Table 1 reports 79 such associations.

There may be many reasons why such variants went unreported, but one reason may be physical proximity to a variant with a stronger signal. Indeed, more than half of the variants described above are within 1Mb of a previously-reported univariate GWAS association. Refraining from reporting multiple near-by associations seems a reasonable – if conservative – strategy to avoid reporting redundant associations due to LD. Further, even when redundant associations due to LD can be ruled out (e.g. by directly examining LD rather than by simply using physical distance), it might be assumed that multiple nearby associated variants may all act through the same biological mechanism and therefore provide redundant biological insights. However, we found that multi-phenotype patterns of association can differ between nearby SNPs, suggesting that they act through different mechanisms.

To highlight just one example, consider rs7515577 – which is an original univariate association in GlobalLipids2010 – and rs12038699 – which is a new multivariate association in GlobalLipids2013. We note that rs12038699 actually reached univariate genome-wide significance in the GlobalLipids2013 dataset, but was not reported (Supplementary Table 6). This is possibly because the latter SNP is relatively close, in genomic terms, to the former SNP (549kb). However, these SNPs are not in strong LD (*r*^2^ = .08), and so these associations almost certainly represent non-redundant associations. This is further supported by the effect sizes in each phenotype, which clearly reveal very different multivariate patterns of effect sizes among phenotypes (Supplementary Figure 2 & Supplementary Table 6). Indeed the very different multivariate patterns of effect size suggest that not only are these associations non-redundant but likely involve different biological mechanisms as well.

These results suggest that, moving forward, it may pay to be more careful in designing filters designed to avoid reporting redundant associations, and that multi-phenotype analyses may have a helpful role to play here.

### Limitations

One goal of the multivariate approach introduced in Stephens 2013 was to increase interpretability of multivariate analyses; in particular, the goal was to not only *test* for associations but also to help *explain* the associations by partitioning the phenotypes into “Unassociated”, “Directly Associated”, and “Indirectly Associated” categories. In principle one might hope to use these classifications to gain insights into the relationships among phenotypes and also perhaps to identify different “types” of multivariate association - effectively clustering associations into different groups. However, in practice we find that these discrete classifications are often not as helpful as one might hope. One reason is the difficulty of reliably distinguishing between direct and indirect effects (Stephens, 2013). Another reason is widespread associations with multiple phenotypes. Indeed, we find that, consistently across data sets, the most common multivariate models involve associations – either direct or indirect – with many phenotypes (Supplementary Table 7) and many SNPs are classified as being associated with many phenotypes (Figure 4A). Further, SNPs are very rarely confidently classified as “Unassociated” with any phenotype (Figure 4B). This last observation can be explained by the fact that it is essentially impossible to distinguish ‘unassociated’ from ‘weakly associated’. Nonetheless when all SNPs show similar classifications it is difficult to get insights into different patterns of multivariate association.

**Figure 4:**
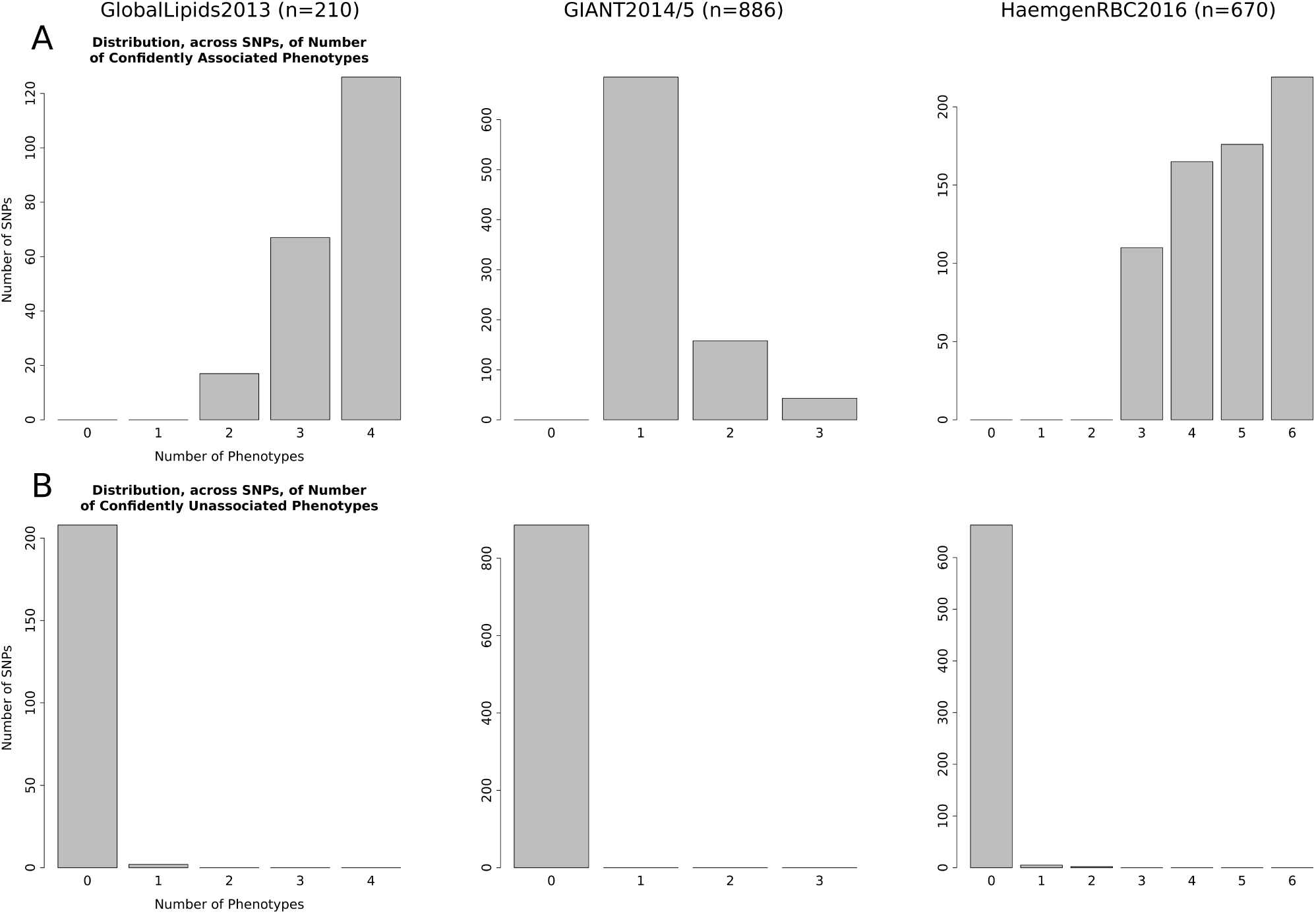
Distribution, Across Significant SNPs, of Number of Phenotypes That Are Confidently Associated (A) or Confidently Unassociated (B). Results are shown for three well-powered datasets: GlobalLipids2013, GIANT2014/5, and HaemgenRBC2016. Here “confident” means with probability *>* 0.95, so a SNP is considered “confidently associated” with a phenotype if the sum of its probabilities in the “Directly Associated” and “Indirectly Associated” categories exceeds 0.95 (A), and is considered confidently unassociated with the phenotype if this probability is less than 0.05 (B). The set of significant SNPs includes both previous univariate associations and new multivariate associations.

Moving forward, rather than relying on the discrete classifications of “Unassociated”, “Directly Associated”, and “Indirectly Associated” to identify different patterns of multivariate association, we believe it will be more fruitful to use multivariate methods that take a more quantitative approach, such as identifying different patterns of *effect size* (including direction of effect) among phenotypes (Urbut et al., 2017). Focusing on effect sizes has the potential to be much more informative than discrete classification, which can hide effect size differences. For example, when multiple SNPs are classified as associated with all phenotypes, they can still show very different patterns of estimated effect sizes/direction (see Supplementary Figure 3).

Another limitation of our multivariate methods is that they can lead to (what appear to be) false positive associations when applied to test SNPs with very low minor allele frequencies. Specifically we saw examples where very low-frequency SNPs (e.g. MAF < .001) showed strong signals of multivariate association despite showing very little signal in any univariate test. Although such results are not impossible, we believe that most of these cases were likely false positives, and we applied a MAF cut-off (of 0.01 or 0.005) to avoid these issues. Consequently we recommend caution in interpreting results of multivariate analyses at very low-frequency SNPs, and more generally we recommend that multivariate results be compared against univariate results to check they make sense – highly significant multivariate associations that do not also show at least a moderate level of univariate association should be treated with caution.

## 4 Discussion

We reanalyzed 13 publicly available GWAS datasets using a Bayesian multivariate approach and identified many new genetic associations. Turning genetic associations into biological discoveries remains, of course, a challenging problem. Nonetheless, our results suggest that the increased power of multivariate association analysis that has been reported in many simulation studies (Stephens, 2013; Galesloot et al., 2014; Porter and O’Reilly, 2017) also translates to discovery of many new associations in practice.

Our results exploit the public availability of summary data from several large GWAS. Despite progress toward easier availability of individual-level data for large studies (Sudlow et al., 2015), in many cases summary data remain much easier to obtain and work with; there are big practical advantages as well to modular pipelines that first compute summary data and then use these as inputs to sub-sequent (more sophisticated) analyses. For example, the multivariate analyses we present here are simplified by assuming that the summary data were computed while adequately adjusting for population stratification. And our results illustrate the potential for reanalysis of summary data to yield novel inferences. In this regard we also emphasize the importance of consortia releasing carefully-chosen summaries. For example, *Z*-scores are much more helpful than *p*-values because they preserve information on the direction of the effect. Even better would be both the effect size and standard error that created the *Z*-score. More generally, although not necessarily essential for our analyses here, it is always helpful to include additional key meta-data (e.g. the reference allele, or effect allele, the minor allele frequency, and sample size).

The specific multivariate methods used here were derived under the assumption that the summary data from each phenotype has been obtained from the same sampled individuals (which is true, at least approximately, for studies analyzed here). However, multivariate analysis of summary data is also possible even when data were obtained from different samples for each phenotype. The main difference between these settings is that, for data from overlapping samples, the “noise” is correlated as well as the signal: i.e. the summary data are correlated under the null due to sample overlap, and correlated under the alternative due to both sample overlap and any shared genetic effects. In contrast, for data from non-overlapping samples the noise is uncorrelated (whereas the signal may remain correlated if genetic factors are shared). Our methods use data at (empirically) null SNPs to estimate the noise correlation, and so their overall assessment of associations should be relatively robust to whether samples for different phenotypes overlap (however, our definitions of **D** (direct) vs **I** (indirect) associations requires the same samples to be measured across phenotypes.)

Moving forward, we expect multivariate association analyses to play an increasingly important role in detecting and understanding genetic associations and relationships among phenotypes. Large studies are now collecting, and making available, rich human genetic and phenotypic information on many complex phenotypes, most notably the UKBioBank (Sudlow et al., 2015). In addition, there are increasingly large studies linking genetic variation and molecular phenotypes such as gene expression (e.g. the GTEx project (GTEx Consortium, 2013)), as well as epigenetic modifications and transcript degradation (Gaffney, 2013; Pai et al., 2015; Birney et al., 2016; Stricker et al., 2017). Analysis of multiple molecular traits can help yield insights into causal connections among traits (Li et al., 2016), and joint analysis of molecular traits with complex phenotypes may also shed light on functional mechanisms (as in “co-localization” analyses (Hormozdiari et al., 2016; Li and Kellis, 2016; Zhu et al., 2016; Wen et al., 2017)). Even simply moving from single phenotype to pairwise analysis can shed considerable light on sharing of genetic effects and possible causal connections (Pickrell et al., 2016; Shi et al., 2017).

These increasingly-complex new data also bring new analytic and computational challenges. Here we have restricted our analyses to relatively small sets of closely-related traits, and indeed the specific multivariate framework we used here – which performs an exhaustive search over all possible multivariate models – is fully tractable for only moderate numbers of traits (up to about 10). Scaling methods up to dealing with larger number of traits may well be helpful for some settings, and recent multivariate analysis methods can deal with dozens of outcomes (Dahl et al., 2016; Urbut et al., 2017). In addition, developing multivariate methods to perform *fine-mapping* of genetic associations simultaneously across multiple phenotypes (Lewin et al., 2016) seems an important and challenging area for future work.

## 5 URLs

bmass R package: https://github.com/mturchin20/bmass

## 6 Acknowledgments

We thank John Novembre, Anna Di Rienzo, and Xin He for helpful feedback during the development of this project. We also thank Peter Carbonetto for helpful feedback on the bmass R package and the manuscript. This work was supported by National Institutes of Health (NIH) Grant R01 HG002585 to MS, NIH Grants T32 GM007197, TL1 TR000432, and F31 AI118375 to MCT, and NIH Grant R01 GM118652.

## 7 Author Contributions

MS conceived the original statistical framework. MS and MCT conceived the study design. MCT performed the data collection, processing, and analyses. MCT wrote the R package bmass. MS supervised the project. MCT and MS wrote the paper.

## 8 Materials and Methods

### 8.1 GWAS Datasets

Below are specific details regarding retrieval and data-processing for each dataset analyzed. Where applicable, these details include the sample size (*N*), minor allele frequency (MAF), and *p*-value thresholds that were applied (based on the thresholds used in the original publications). For each dataset variants were dropped if they satisfied at least one of the following criteria: did not contain information for every phenotype; had missing MAF; were fixed (MAF of 0); had effect size exactly 0 (i.e. direction of effect would be indeterminable); or did not contain the same reference and alternative alleles across each phenotype. For a handful of studies, external databases were used to retrieve chromosome, basepair information, and MAF based on rsID#; in these studies SNPs for which this information could not be retrieved were also dropped.

**GlobalLipids2010 (Teslovich et al., 2010):** Original merged, processed, and GWAS-hit annotated summary data from Stephens 2013 (Stephens, 2013) for HDL, LDL, TG, and TC was downloaded from https://github.com/stephens999/multivariate (*dtlesssignif.annot.txt* and *RSS0.txt*).

**GlobalLipids2013 (Willer et al., 2013):** Summary data for HDL, LDL, TG, and TC was downloaded from http://csg.sph.umich.edu/abecasis/public/lipids2013/. We used a minimum *N* threshold of 50,000, a MAF threshold of 1%, and a univariate significant GWAS *p*-value threshold of 5 *×* 10*^−^*^8^. All variants were oriented to the HDL minor allele. The final merged and QC’d datafile contained 2,004,701 SNPs. rsID#’s of published GWAS SNPs were retrieved for all four phenotypes from https://www.nature.com/ng/journal/v45/n11/full/ng.2797.html via Supplementary Tables 2 and 3.

**GIANT2010 (Lango Allen et al., 2010; Speliotes et al., 2010; Heid et al., 2010):** Summary data for Height, BMI, and WHRadjBMI were downloaded from https://www.broadinstitute.org/collaboration/giant/index. php/GIANT_consortium_data_files. We used a minimum *N* threshold of 50,000, a MAF threshold of 1%, and a univariate significant GWAS *p*-value threshold of 5 *×* 10*^−^*^8^. Chromosome and basepair position per variant were retrieved from dbSNP130 (Sherry et al., 2001). All variants were oriented to the Height minor allele. The final merged and QC’ed datafile contained 2,363,881 SNPs. rsID#’s of published GWAS SNPs were retrieved for Height from https://www.nature.com/nature/journal/v467/n7317/full/nature09410.html via Supplementary Table 1, for BMI from https://www.nature.com/ng/journal/v42/n11/full/ng.686.html via Table 1, and for WHRadjBMI from https://www.nature.com/ng/journal/v42/n11/full/ng.685.html via Table 1.

**GIANT2014/5 (Wood et al., 2014; Locke et al., 2015; Shungin et al., 2015):** Summary data for Height, BMI, and WHRadjBMI were downloaded from https://www.broadinstitute.org/collaboration/giant/index.php/GIANT_consortium_data_files. We used a minimum *N* threshold of 50,000, a MAF threshold of 1%, and a univariate significant GWAS *p*-value threshold of 5 *×* 10*^−^*^8^. Chromosome and basepair position per variant were retrieved from dbSNP130 (Sherry et al., 2001). All variants were oriented to the Height minor allele. The final merged and QC’ed datafile contained 2,340,715 SNPs. rsID#’s of published GWAS SNPs were retrieved for Height from https://www.nature.com/ng/journal/v46/n11/full/ng.3097.html via Supplementary Table 1, for BMI from https://www.nature.com/nature/journal/v518/n7538/full/nature14177.html via Supplementary Tables 1 and 2, and for WHRadjBMI from https://www.nature.com/nature/journal/v518/n7538/full/nature14132.html via Supplementary Table 4.

**HaemgenRBC2012 (van der Harst et al., 2012):** Summary data for RBC, PCV, MCV, MCH, MCHC, and Hb were downloaded from the European Genome-Phenome Archive via accession number EGAS00000000132 (https://www.ebi.ac.uk/ega/studies/EGAS00000000132). We used a minimum *N* threshold of 10,000, a MAF threshold of 1%, and a univariate significant GWAS *p*-value threshold of 1 *×* 10*^−^*^8^. Chromosome, basepair position, and MAF per variant were retrieved from HapMap release 22 (International HapMap, 2003). All variants were oriented to the RBC minor allele. The final merged and QC’ed datafile contained 2,327,567 SNPs. rsID#’s of published GWAS SNPs were retrieved for all six phenotypes from https://www.nature.com/nature/journal/v492/n7429/full/nature11677.html via Table 1.

**HaemgenRBC2016 (Astle et al., 2016):** Summary data for RBC, PCV, MCV, MCH, MCHC, and Hb were shared via personal communication with the authors. We used a MAF threshold of 1% and a univariate significant GWAS *p*-value threshold of 8.319*×*10*^−^*^9^. Since sample size was not provided per variant, the following overall study sample sizes were used as proxies per phenotype: 172,952 for RBC, 172,433 for PCV, 173,039 for MCV, 172,332 for MCH, for 172,925 MCHC, and 172,851 for Hb. All variants were oriented to the RBC minor allele. Only SNPs were analyzed. The final merged and QC’ed datafile contained 8,649,095 SNPs. We then used these summary data to create a list of (non-redundant) “Previous univariate associations”. This was done separately for each phenotype by collecting all SNPs that exceeded the univariate significant GWAS *p*-value threshold and greedily pruning the SNPs: i.e. we went down the list, removing SNPs that were less significant than another SNP within 500kb. The pruned lists of previous univariate associations for each phenotype were then combined to produce the final SNP list of “published GWAS results”. Published CNVs that tagged regions that were not identified by this ‘final published SNP list’ were also included to avoid erroneously claiming downstream a region as a ‘new unpublished result’; these CNVs however were only used to mask additional loci as being ‘nearby a published univariate GWAS result’ and for nothing else in the bmass analysis pipeline.

**ICBP2011 (International Consortium for Blood Pressure Genome-Wide Association et al., 2011; Wain et al., 2011):** Summary data for SBP, DBP, PP, and MAP were downloaded from dbGaP via accession number phs000585.v1.p1 (https://www.ncbi.nlm.nih.gov/projects/gap/cgi-bin/study.cgi?study_id=phs000585.v1.p1). We used a minimum *N* threshold of 10,000, a MAF threshold of 1%, and a univariate significant GWAS *p*-value threshold of 5 *×* 10*^−^*^8^. Chromosome and basepair position per variant were retrieved from HapMap release 21 (International HapMap, 2003). All variants were oriented to the SBP minor allele. The final merged and QC’ed datafile contained 2,387,851 SNPs. rsID#’s of published GWAS SNPs were retrieved for SBP and DBP from https://www.nature.com/nature/journal/v478/n7367/full/nature10405.html via Supplementary Table 5, and for PP and MAP from https://www.nature.com/ng/journal/v43/n10/full/ng.922.html via Table 1 and Supplementary Table 2F. Additionally, we gratefully acknowledge the International Consortium for Blood Pressure Genome-Wide Association Studies (Nature. 2011 Sep 11;478(7367):103-9, Nat Genet. 2011 Sep 11;43(10):1005-11) for generating and sharing these data.

**MAGIC2010 (Dupuis et al., 2010):** Summary data for FstIns, FstGlu, HOMA_B, and HOMA_IR were downloaded from https://www.magicinvestigators.org/downloads/. We used a MAF threshold of 1% and a univariate significant GWAS *p*-value threshold of 5 *×* 10*^−^*^8^. Since sample size was not provided per variant, the overall study sample size of 46,186 was used as a proxy. Chromosome and basepair position per variant were retrieved from HapMap release 22 (International HapMap, 2003). All variants were oriented to the FstIns minor allele. The final merged and QC’ed datafile contained 2,333,328 SNPs. rsID#’s of published GWAS SNPs were retrieved for all four phenotypes from https://www.nature.com/ng/journal/v42/n2/full/ng.520.html via Table 1.

**GEFOS2015 (Zheng et al., 2015):** Summary data for FA, FN, and LS were downloaded from http://www.gefos.org/?q=content/data-release-2015. We used a MAF threshold of .5% and a univariate significant GWAS *p*-value threshold of 1.2*×*10*^−^*^8^. Since sample size was not provided per variant, the overall study sample size of 32,965 was used as a proxy. All variants were oriented to the FA minor allele. The final merged and QC’ed datafile contained 8,938,035 SNPs. rsID#’s of published GWAS SNPs were retrieved for all four phenotypes from https://www.nature.com/nature/journal/v526/n7571/full/nature14878.html via Supplementary Table 13.

**GIS2014 (Benyamin et al., 2014):** Summary data for Iron, Sat, TrnsFrn, and Log10Frtn were shared via personal communication with the authors. We used a MAF threshold of 1% and a univariate significant GWAS *p*-value threshold of 5 *×* 10*^−^*^8^. Since sample size was not provided per variant, the overall study sample size of 48,972 was used as a proxy. All variants were oriented to the Iron minor allele. The final merged and QC’ed datafile contained 1,985,313 SNPs. rsID#’s of published GWAS SNPs were retrieved for all four phenotypes from https://www.nature.com/articles/ncomms5926/ via Table 1.

**SSGAC2016 (Barban et al., 2016):** Summary data for NEB_Pooled and AFB_Pooled were downloaded from https://www.thessgac.org/data. We used a MAF threshold of 1% and a univariate significant GWAS *p*-value threshold of 5 *×* 10*^−^*^8^. Since sample size was not provided per variant, the following overall study sample sizes were used as proxies per phenotype: 251,151 for NEB_Pooled and 343,072 for AFB_Pooled. All variants were oriented to the NEB_Pooled minor allele. The final merged and QC’ed datafile contained 2,395,561 SNPs. rsID#’s of published GWAS SNPs were retrieved for all four phenotypes from https://www.nature.com/ng/journal/v48/n12/full/ng.3698.html via Table 1.

**CKDGen2010/1 (Kottgen et al., 2010; Boger et al., 2011):** Summary data for Crea, Cys, CKD, UACR, and MA were downloaded from https://www.nhlbi.nih.gov/research/intramural/researchers/pi/fox-caroline/datasets. We used a MAF threshold of 1% and a univariate significant GWAS *p*-value thresh-old of 5 *×* 10*^−^*^8^. Since sample size was not provided per variant, the following overall study sample sizes were used as proxies per phenotype: 67,093 for Crea, 20,957 for Cys, 62,237 for CKD, 31,580 for UACR, and 30,482 for MA. All variants were oriented to the Crea minor allele. The final merged and QC’ed datafile contained 2,333,498 SNPs. rsID#’s of published GWAS SNPs were retrieved for Crea, Cys, and CKD from https://www.nature.com/ng/journal/v42/n5/full/ng.568.html via Table 2.

**Table 2:** Summary of New Multivariate Associations Identified. Previous Univariate: the number of previous genome-wide significant univariate associations based on the publicly available summary data. New Multivariate: the number of new genome-wide significant multivariate associations. BF_av_ Thresh: the Bayes Factor threshold used in declaring new multivariate associations to be significant. Overlap With Next Release: for GlobalLipids2010, GIANT2010, and HaemgenRBC2012, the last column shows the number of new multivariate associations that overlap with the univariate GWAS associations in the next release from the same consortium; overlap is defined as being within 50kb of the univariate GWAS variant.

| Dataset | Release | —SNP Associations— | | $\text{BF}_{\text{av}}$<br>Thresh | Overlap With<br>Next Release |
| --- | --- | --- | --- | --- | --- |
|  |  | Previous<br>Univariate | New<br>Multivariate |  |  |
| GlobalLipids | 2010 | 102 | 19 | 4.35 | 13/19 |
|  | 2013 | 145 | 65 | 4.29 | - |
| GIANT | 2010 | 144 | 60 | 4.11 | 49/60 |
|  | 2014/5 | 724 | 162 | 4.49 | - |
| HaemgenRBC | 2012 | 63 | 16 | 5.21 | 9/16 |
|  | 2016 | 610 | 60 | 4.68 | - |
| ICBP | 2011 | 22 | 22 | 5.24 | - |
| MAGIC | 2010 | 12 | 1 | 6.90 | - |
| GEFOS | 2015 | 34 | 13 | 5.06 | - |
| GIS | 2014 | 8 | 5 | 7.04 | - |
| SSGAC | 2016 | 9 | 1 | 5.43 | - |
| CKDGen | 2010/1 | 28 | 6 | 4.10 | - |
| ENIGMA2 | 2015 | 5 | 3 | 7.48 | - |

**ENIGMA22015 (Hibar et al., 2015):** Summary data for ICV, Accumbens, Amygdala, Caudate, Hippocampus, Pallidum, Putamen, and Thalamus were downloaded from http://enigma.ini.usc.edu/research/download-enigma-gwas-results/. We used a minimum *N* threshold of 10,000, a MAF threshold of 1% and a univariate significant GWAS *p*-value threshold of 5 *×* 10*^−^*^8^. All variants were oriented to the ICV minor allele. The final merged and QC’ed datafile contained 6,271,117 SNPs. rsID#’s of published GWAS SNPs were retrieved for all 8 phenotypes from https://www.nature.com/nature/journal/v520/n7546/full/nature14101.html via Table 1.

### 8.2 bmass

bmass implements in an R package the statistical methods described in Stephens 2013, which should be consulted for full details. In particular, the sections “Computation” and “Detailed Methods (Global Lipids Analysis)” in Stephens 2013 describe how multivariate analyses are applied to GWAS summary data, and bmass implements the data analysis pipeline described in the “Detailed Methods (Global Lipids Analysis)” section. The bmass R package also includes two vignettes to help users begin processing GWAS summary data and implementing these methods.

### 8.3 Additional Details for Figure 3

For each dataset we made a list of “marginally-significant” SNPs, with *p*-values smaller than 1 *×* 10*^−^*^6^ but not genome-wide significant at the relevant datasets’ GWAS threshold. We then greedily pruned these lists of marginally-significant SNPs: that is we repeatedly went through the lists removing SNPs that were less significant than another SNP within 500kb. We then removed any SNPs that were within 500kb of a new multivariate association, and merged the resulting list with the list of new multivariate associations, and sorted this merged list of SNPs by their minimum univariate *p*-value.

This results in a non-redundant list of marginally-significant SNPs – some of which are new multivariate associations and some of which are not – sorted by their smallest univariate *p*-value. The plot shows how the number of SNPs of each type varies as the *p*-value threshold is relaxed from the GWAS threshold to 10*^−^*^6^ (the HaemgenRBC2016 results show only the top 500 SNPs due to the abundance of SNPs between 8.31 *×* 10*^−^*^9^ and 1 *×* 10*^−^*^6^).

## 9 Supporting Information Legends

Supplementary Figure 1: **Graphical Model of Multivariate Categories**. Shown here is a Directed Acyclic Graphical (DAG) model of our multivariate categories in the context of our vector of phenotypes **Y** (e.g. **Y** = {**Y_U_**, **Y_D_**, **Y_I_**}) and their connections with the variant of interest **g**. The relationships described in-text can be seen here. **Y_U_**, our unassociated phenotypes, have no connection with **g**. **Y_D_**, our directly associated phenotypes, have a direct connection with **g**. And **Y_I_**, our indirectly associated phenotypes, have a connection with **g** only by going through **Y_D_** first. Note that if **Y_D_** were not observed, **Y_I_** would appear as a direct connection.

Supplementary Figure 2: **Refining Association Signals – GlobalLipids2013 rs7515577 & rs12038699**. Shown are the -log_10_ univariate *p*-values from the GlobalLipids2013 analysis for both the previous univariate association rs7515577 (“Previous Univariate SNP”) and the new multivariate association rs12038699 (“New Multivariate SNP”) across all four phenotypes analyzed. rs7515577 is represented as a triangle and rs12038699 is represented as a square. Also shown are the -log_10_ univariate *p*-values of SNPs within 1Mb of the midpoint between rs7515577 and rs12038699. Color-coding of the SNPs represent the degree of linkage disequilibrium between variants and the new association rs12038699 based on the GBR cohort of 1000Genomes (Genomes Project et al., 2015); for color coding details, see legend.

Supplementary Figure 3: **Effect Size Heterogeneity Among SNPs With Identical Multivariate Model Assignments**. Shown are the phenotype effect sizes (points), and *±*2 standard errors (bars), for four significantly associated SNPs from HaemgenRBC2016. All four SNPs were classified as being “associated” with all six phenotypes (i.e. marginal posterior probability of association >= 95% for each phenotype). However, they clearly show different patterns of effect sizes. Therefore focusing simply on binary calls of “associated” vs “unassociated” can hide different patterns of multivariate association.

Supplementary Table 1: Summary of Associations in Each Dataset.

^a^Number of new multivariate associations discovered by our analysis. Note that we required a multivariate association to be at least 500kb from a previous reported association to be considered “new”.

^b^Univariate GWAS significance *p*-value threshold used by the original study publication.

^c^These are new multivariate SNPs that were not reported by the original study despite having a univariate association (in the public summary data) that was genome-wide significant by the original study’s univariate significance threshold.

^d^A “previous association” means an association reported by the original GWAS; “near” means within 1Mb (but these are all more than 500kb away from a previous association since our classification of new multivariate SNPs requires this).

Supplementary Tables 2a-m: **Lists of New** bmass **Multivariate Associations, per Dataset**. Attached Excel sheets list new bmass associations for each dataset analyzed.

Supplementary Tables 3a-m: **Lists of Retrieved Univariate Associations From Original Publications, per Dataset**. Attached Excel sheets list the rsID#’s of the univariate significant SNPs that were retrieved from the original publication(s) associated with each dataset (see Online Methods for details).

Supplementary Tables 4a-m: **Results for Previous Univariate Associations, per Dataset**. Attached Excel sheets give bmass results for previous univariate associations, per dataset. Note that these results may not include all SNPs from Tables 3a-m, because some SNPs were dropped during QC and other SNPs were dropped because they did not reach univariate significance in the publicly available summary data (see Online Methods for details).

Supplementary Table 5: **Replication of New Multivariate Associations**. Shown are example metrics of how well our new multivariate associations replicate in datasets that allow such an evaluation. Specifically, for three of the studies used (GlobalLipids, GIANT, and HaemgenRBC), there are multiple dataset releases. To examine how well our new multivariate bmass associations replicate, we compared the results from the first releases (“1^st^”) with the univariate GWAS associations of the second releases (“2^nd^”). In essence, each of these approaches aim to increase power – one by using a multivariate approach (bmass) and the other by increasing sample size (the 2^nd^ releases) – thus allowing us to compare the results against one another. Univariate *p*-Value Threshold: univariate GWAS significance *p*-value thresholds used by the original publication(s) for both the earlier (1^st^) and later (2^nd^) releases. New Multivariate SNPs in 1^st^: number of new multivariate associations from the earlier release. Lower Univariate *p*-Value in 2^nd^: number of new multivariate associations from the earlier release that also have lower univariate *p*-values in the later release. Below 2^nd^ Univariate Threshold: number of new multivariate associations from the earlier release that also cross the later release’s univariate GWAS significance threshold.

Supplementary Table 6: *p***-Values for rs7515577 & rs12038699 in 2010 and 2013 GlobalLipds Releases** – In the 2010 release rs7515577 has a univariate *p*-value that crosses the 5 *×* 10*^−^*^8^ threshold (TC), whereas rs12038699 does not. Since rs12038699 is near to rs7515577 it may get masked for future analyses; however in the 2013 data rs12038699 not only has a lower minimum univariate *p*-value, but also has a different multivariate *p*-value pattern as compared to rs7515577. Both these signals suggest that rs12038699 should be viewed as a separate GWAS hit for GlobalLipids2013.

Supplementary Table 7: **Top Multivariate Model Examples per SNP**. List of multivariate models that most frequently have the highest posterior probabilities per SNP. Top 5 models are shown from across both the previous univariate associations analyzed and the new multivariate associations discovered in the GlobalLipids2013, GIANT2014/5, and HaemgenRBC2016 datasets. Phenotype ordering is shown in the header, where 0, 1, and 2 refer to the multivariate categories of **U**nassociated, **D**irectly Associated, and **I**ndirectly Associated. n is the number of SNPs that show the specified model as having the largest posterior probability, with Mean Posterior displaying the average posterior probability of the given model across the n SNPs, and Original Prior showing the prior established for the given model from training on all the previous univariate associations from that dataset.

## References

1. Astle, W. J., Elding, H., Jiang, T., Allen, D., Ruklisa, D., Mann, A. L., Mead, D., Bouman, H., Riveros-Mckay, F., Kostadima, M. A., Lambourne, J. J., Sivapalaratnam, S., Downes, K., Kundu, K., Bomba, L., Berentsen, K., Bradley, J. R., Daugherty, L. C., Delaneau, O., Freson, K., Garner, S. F., Grassi, L., Guerrero, J., Haimel, M., Janssen-Megens, E. M., Kaan, A., Kamat, M., Kim, B., Mandoli, A., Marchini, J., Martens, J. H., Meacham, S., Megy, K., O’Connell, J., Petersen, R., Sharifi, N., Sheard, S. M., Staley, J. R., Tuna, S., van der Ent, M., Walter, K., Wang, S. Y., Wheeler, E., Wilder, S. P., Iotchkova, V., Moore, C., Sambrook, J., Stunnenberg, H. G., Di Angelantonio, E., Kaptoge, S., Kuijpers, T. W., Carrillo-de Santa-Pau, E., Juan, D., Rico, D., Valencia, A., Chen, L., Ge, B., Vasquez, L., Kwan, T., Garrido-Martin, D., Watt, S., Yang, Y., Guigo, R., Beck, S., Paul, D. S., Pastinen, T., Bujold, D., Bourque, G., Frontini, M., Danesh, J., Roberts, D. J., Ouwehand, W. H., Butterworth, A. S., and Soranzo, N. (2016). The allelic landscape of human blood cell trait variation and links to common complex disease. Cell, 167(5):1415–1429 e19.

2. Barban, N., Jansen, R., de Vlaming, R., Vaez, A., Mandemakers, J. J., Tropf, F. C., Shen, X., Wilson, J. F., Chasman, D. I., Nolte, I. M., Tragante, V., van der Laan, S. W., Perry, J. R., Kong, A., Consortium, B., Ahluwalia, T. S., Albrecht, E., Yerges-Armstrong, L., Atzmon, G., Auro, K., Ayers, K., Bakshi, A., Ben-Avraham, D., Berger, K., Bergman, A., Bertram, L., Bielak, L. F., Bjornsdottir, G., Bonder, M. J., Broer, L., Bui, M., Barbieri, C., Cavadino, A., Chavarro, J. E., Turman, C., Concas, M. P., Cordell, H. J., Davies, G., Eibich, P., Eriksson, N., Esko, T., Eriksson, J., Falahi, F., Felix, J. F., Fontana, M. A., Franke, L., Gandin, I., Gaskins, A. J., Gieger, C., Gunderson, E. P., Guo, X., Hayward, C., He, C., Hofer, E., Huang, H., Joshi, P. K., Kanoni, S., Karlsson, R., Kiechl, S., Kifley, A., Kluttig, A., Kraft, P., Lagou, V., Lecoeur, C., Lahti, J., Li-Gao, R., Lind, P. A., Liu, T., Makalic, E., Mamasoula, C., Matteson, L., Mbarek, H., McArdle, P. F., McMahon, G., Meddens, S. F., Mihailov, E., Miller, M., Missmer, S. A., Monnereau, C., van der Most, P. J., Myhre, R., Nalls, M. A., Nutile, T., Kalafati, I. P., Porcu, E., Prokopenko, I., Rajan, K. B., Rich-Edwards, J., Rietveld, C. A., Robino, A., Rose, L. M., Rueedi, R., Ryan, K. A., Saba, Y., Schmidt, D., Smith, J. A., Stolk, L., Streeten, E., Tonjes, A., Thorleifsson, G., et al. (2016). Genome-wide analysis identifies 12 loci influencing human reproductive behavior. Nat Genet, 48(12):1462–1472.

3. Benyamin, B., Esko, T., Ried, J. S., Radhakrishnan, A., Vermeulen, S. H., Traglia, M., Gogele, M., Anderson, D., Broer, L., Podmore, C., Luan, J., Kutalik, Z., Sanna, S., van der Meer, P., Tanaka, T., Wang, F., Westra, H. J., Franke, L., Mihailov, E., Milani, L., Halldin, J., Winkelmann, J., Meitinger, T., Thiery, J., Peters, A., Waldenberger, M., Rendon, A., Jolley, J., Sambrook, J., Kiemeney, L. A., Sweep, F. C., Sala, C. F., Schwienbacher, C., Pichler, I., Hui, J., Demirkan, A., Isaacs, A., Amin, N., Steri, M., Waeber, G., Verweij, N., Powell, J. E., Nyholt, D. R., Heath, A. C., Madden, P. A., Visscher, P. M., Wright, M. J., Montgomery, G. W., Martin, N. G., Hernandez, D., Bandinelli, S., van der Harst, P., Uda, M., Vollenweider, P., Scott, R. A., Langenberg, C., Wareham, N. J., InterAct, C., van Duijn, C., Beilby, J., Pramstaller, P. P., Hicks, A. A., Ouwehand, W. H., Oexle, K., Gieger, C., Metspalu, A., Camaschella, C., Toniolo, D., Swinkels, D. W., and Whitfield, J. B. (2014). Novel loci affecting iron homeostasis and their effects in individuals at risk for hemochromatosis. Nat Commun, 5:4926.

4. Birney, E., Smith, G. D., and Greally, J. M. (2016). Epigenome-wide association studies and the interpretation of disease -omics. PLoS Genet, 12(6):e1006105.

5. Boger, C. A., Chen, M. H., Tin, A., Olden, M., Kottgen, A., de Boer, I. H., Fuchs-berger, C., O’Seaghdha, C. M., Pattaro, C., Teumer, A., Liu, C. T., Glazer, N. L., Li, M., O’Connell, J. R., Tanaka, T., Peralta, C. A., Kutalik, Z., Luan, J., Zhao, J. H., Hwang, S. J., Akylbekova, E., Kramer, H., van der Harst, P., Smith, A. V., Lohman, K., de Andrade, M., Hayward, C., Kollerits, B., Tonjes, A., Aspelund, T., Ingelsson, E., Eiriksdottir, G., Launer, L. J., Harris, T. B., Shuldiner, A. R., Mitchell, B. D., Arking, D. E., Franceschini, N., Boerwinkle, E., Egan, J., Hernandez, D., Reilly, M., Townsend, R. R., Lumley, T., Siscovick, D. S., Psaty, B. M., Kestenbaum, B., Haritunians, T., Bergmann, S., Vollenweider, P., Waeber, G., Mooser, V., Waterworth, D., Johnson, A. D., Florez, J.C., Meigs, J. B., Lu, X., Turner, S. T., Atkinson, E. J., Leak, T. S., Aasarod, K, Skorpen, F., Syvanen, A. C., Illig, T., Baumert, J., Koenig, W., Kramer, B. K., Devuyst, O., Mychaleckyj, J. C., Minelli, C., Bakker, S. J., Kedenko, L., Paulweber, B., Coassin, S., Endlich, K., Kroemer, H. K., Biffar, R., Stracke, S., Volzke, H., Stumvoll, M., Magi, R., Campbell, H., Vitart, V., Hastie, N. D., Gudnason, V., Kardia, S. L., Liu, Y., Polasek, O., Curhan, G., Kronenberg, F., Prokopenko, I., Rudan, I., Arnlov, J., Hallan, S., Navis, G., Consortium, C. K., Parsa, A., Ferrucci, L., Coresh, J., Shlipak, M. G., et al. (2011). Cubn is a gene locus for albuminuria. J Am Soc Nephrol, 22(3):555–70.

6. Dahl, A., Iotchkova, V., Baud, A., Johansson, A., Gyllensten, U., Soranzo, N., Mott, R., Kranis, A., and Marchini, J. (2016). A multiple-phenotype imputation method for genetic studies. Nat Genet, 48(4):466–72.

7. Dupuis, J., Langenberg, C., Prokopenko, I., Saxena, R., Soranzo, N., Jackson, A. U., Wheeler, E., Glazer, N. L., Bouatia-Naji, N., Gloyn, A. L., Lindgren, C. M., Magi, R., Morris, A. P., Randall, J., Johnson, T., Elliott, P., Rybin, D., Thorleifsson, G., Steinthorsdottir, V., Henneman, P., Grallert, H., Dehghan, A., Hottenga, J. J., Franklin, C. S., Navarro, P., Song, K., Goel, A., Perry, J. R., Egan, J. M., Lajunen, T., Grarup, N., Sparso, T., Doney, A., Voight, B. F., Stringham, H. M., Li, M., Kanoni, S., Shrader, P., Cavalcanti-Proenca, C., Kumari, M., Qi, L., Timpson, N. J., Gieger, C., Zabena, C., Rocheleau, G., Ingelsson, E., An, P., O’Connell, J., Luan, J., Elliott, A., McCarroll, S. A., Payne, F., Roccasecca, R. M., Pattou, F., Sethupathy, P., Ardlie, K., Ariyurek, Y., Balkau, B., Barter, P., Beilby, J. P., Ben-Shlomo, Y., Benediktsson, R., Bennett, A. J., Bergmann, S., Bochud, M., Boerwinkle, E., Bonnefond, A., Bonnycastle, L. L., Borch-Johnsen, K., Bottcher, Y., Brunner, E., Bumpstead, S. J., Charpentier, G., Chen, Y. D., Chines, P., Clarke, R., Coin, L. J., Cooper, M. N., Cornelis, M., Crawford, G., Crisponi, L., Day, I. N., de Geus, E. J., Delplanque, J., Dina, C., Erdos, M. R., Fedson, A. C., Fischer-Rosinsky, A., Forouhi, N. G., Fox, C. S., Frants, R., Franzosi, M. G., Galan, P., Goodarzi, M. O., Graessler, J., Groves, C. J., Grundy, S., Gwilliam, R., Gyllensten, U., Hadjadj, S., et al. (2010). New genetic loci implicated in fasting glucose homeostasis and their impact on type 2 diabetes risk. Nat Genet, 42(2):105–16.

8. Gaffney, D. J. (2013). Global properties and functional complexity of human gene regulatory variation. PLoS Genet, 9(5):e1003501.

9. Galesloot, T. E., van Steen, K., Kiemeney, L. A., Janss, L. L., and Vermeulen, S. H. (2014). A comparison of multivariate genome-wide association methods. PLoS One, 9(4):e95923.

10. Genomes Project, C., Auton, A., Brooks, L. D., Durbin, R. M., Garrison, E. P., Kang, H. M., Korbel, J. O., Marchini, J. L., McCarthy, S., McVean, G. A., and Abecasis, G. R. (2015). A global reference for human genetic variation. Nature, 526(7571):68–74.

11. GTEx Consortium, T. (2013). The genotype-tissue expression (gtex) project. Nat Genet, 45(6):580–5.

12. Heid, I. M., Jackson, A. U., Randall, J. C., Winkler, T. W., Qi, L., Steinthorsdottir, V., Thorleifsson, G., Zillikens, M. C., Speliotes, E. K., Magi, R., Workalemahu, T., White, C. C., Bouatia-Naji, N., Harris, T. B., Berndt, S. I., Ingelsson, E., Willer, C. J., Weedon, M. N., Luan, J., Vedantam, S., Esko, T., Kilpelainen, T. O., Kutalik, Z., Li, S., Monda, K. L., Dixon, A. L., Holmes, C. C., Kaplan, L. M., Liang, L., Min, J. L., Moffatt, M. F., Molony, C., Nicholson, G., Schadt, E. E., Zondervan, K. T., Feitosa, M. F., Ferreira, T., Lango Allen, H., Weyant, R. J., Wheeler, E., Wood, A. R., Magic, Estrada, K., Goddard, M. E., Lettre, G., Mangino, M., Nyholt, D. R., Purcell, S., Smith, A. V., Visscher, P. M., Yang, J., McCarroll, S. A., Nemesh, J., Voight, B. F., Absher, D., Amin, N., Aspelund, T., Coin, L., Glazer, N. L., Hayward, C., Heard-Costa, N. L., Hottenga, J. J., Johansson, A., Johnson, T., Kaakinen, M., Kapur, K., Ketkar, S., Knowles, J. W., Kraft, P., Kraja, A. T., Lamina, C., Leitzmann, M. F., McKnight, B., Morris, A. P., Ong, K. K., Perry, J. R., Peters, M. J., Polasek, O., Prokopenko, I., Rayner, N. W., Ripatti, S., Rivadeneira, F., Robertson, N. R., Sanna, S., Sovio, U., Surakka, I., Teumer, A., van Wingerden, S., Vitart, V., Zhao, J. H., Cavalcanti-Proenca, C., Chines, P. S., Fisher, E., Kulzer, J. R., Lecoeur, C., Narisu, N., Sandholt, C., Scott, L. J., Silander, K., Stark, K., et al. (2010). Meta-analysis identifies 13 new loci associated with waist-hip ratio and reveals sexual dimorphism in the genetic basis of fat distribution. Nat Genet, 42(11):949–60.

13. Hibar, D. P., Stein, J. L., Renteria, M. E., Arias-Vasquez, A., Desrivieres, S., Jahanshad, N., Toro, R., Wittfeld, K., Abramovic, L., Andersson, M., Aribisala, B. S., Armstrong, N. J., Bernard, M., Bohlken, M. M., Boks, M. P., Bralten, J., Brown, A. A., Chakravarty, M. M., Chen, Q., Ching, C. R., Cuellar-Partida, G., den Braber, A., Giddaluru, S., Goldman, A. L., Grimm, O., Guadalupe, T., Hass, J., Woldehawariat, G., Holmes, A. J., Hoogman, M., Janowitz, D., Jia, T., Kim, S., Klein, M., Kraemer, B., Lee, P. H., Olde Loohuis, L. M., Luciano, M., Macare, C., Mather, K. A., Mattheisen, M., Milaneschi, Y., Nho, K., Papmeyer, M., Ramasamy, A., Risacher, S. L., Roiz-Santianez, R., Rose, E. J., Salami, A., Samann, P. G., Schmaal, L., Schork, A. J., Shin, J., Strike, L. T., Teumer, A., van Donkelaar, M. M., van Eijk, K. R., Walters, R. K., Westlye, L. T., Whelan, C. D., Winkler, A. M., Zwiers, M. P., Alhusaini, S., Athanasiu, L., Ehrlich, S., Hakobjan, M. M., Hartberg, C. B., Haukvik, U. K., Heister, A. J., Hoehn, D., Kasperaviciute, D., Liewald, D. C., Lopez, L. M., Makkinje, R. R., Matarin, M., Naber, M. A., McKay, D. R., Needham, M., Nugent, A. C., Putz, B., Royle, N. A., Shen, L., Sprooten, E., Trabzuni, D., van der Marel, S. S., van Hulzen, K. J., Walton, E., Wolf, C., Almasy, L., Ames, D., Arepalli, S., Assareh, A. A., Bastin, M. E., Brodaty, H., Bulayeva, K. B., Carless, M. A., Cichon, S., Corvin, A., Curran, J. E., Czisch, M., et al. (2015). Common genetic variants influence human subcortical brain structures. Nature, 520(7546):224–9.

14. Hormozdiari, F., van de Bunt, M., Segre, A. V., Li, X., Joo, J. W. J., Bilow, M., Sul, J. H., Sankararaman, S., Pasaniuc, B., and Eskin, E. (2016). Colocalization of gwas and eqtl signals detects target genes. Am J Hum Genet, 99(6):1245–1260.

15. International Consortium for Blood Pressure Genome-Wide Association, S., Ehret, G. B., Munroe, P. B., Rice, K. M., Bochud, M., Johnson, A. D., Chasman, D. I., Smith, A. V., Tobin, M. D., Verwoert, G. C., Hwang, S. J., Pihur, V., Vollenweider, P., O’Reilly, P. F., Amin, N., Bragg-Gresham, J. L., Teumer, A., Glazer, N. L., Launer, L., Zhao, J. H., Aulchenko, Y., Heath, S., Sober, S., Parsa, A., Luan, J., Arora, P., Dehghan, A., Zhang, F., Lucas, G., Hicks, A. A., Jackson, A. U., Peden, J. F., Tanaka, T., Wild, S. H., Rudan, I., Igl, W., Milaneschi, Y., Parker, A. N., Fava, C., Chambers, J. C., Fox, E. R., Kumari, M., Go, M. J., van der Harst, P., Kao, W. H., Sjogren, M., Vinay, D. G., Alexander, M., Tabara, Y., Shaw-Hawkins, S., Whincup, P. H., Liu, Y., Shi, G., Kuusisto, J., Tayo, B., Seielstad, M., Sim, X., Nguyen, K. D., Lehtimaki, T., Matullo, G., Wu, Y., Gaunt, T. R., Onland-Moret, N. C., Cooper, M. N., Platou, C. G., Org, E., Hardy, R., Dahgam, S., Palmen, J., Vitart, V., Braund, P. S., Kuznetsova, T., Uiterwaal, C. S., Adeyemo, A., Palmas, W., Campbell, H., Ludwig, B., Tomaszewski, M., Tzoulaki, I., Palmer, N. D., consortium, C. A., Consortium, C. K., KidneyGen, C., EchoGen, c., consortium, C.-H., Aspelund, T., Garcia, M., Chang, Y. P., O’Connell, J. R., Steinle, N. I., Grobbee, D. E., Arking, D. E., Kardia, S. L., Morrison, A. C., Hernandez, D., Najjar, S., McArdle, W. L., Hadley, D., Brown, M. J., Connell, J. M., et al. (2011). Genetic variants in novel pathways influence blood pressure and cardiovascular disease risk. Nature, 478(7367):103–9.

16. International HapMap, C. (2003). The international hapmap project. Nature, 426(6968):789–96.

17. Jiang, C. and Zeng, Z. B. (1995). Multiple trait analysis of genetic mapping for quantitative trait loci. Genetics, 140(3):1111–27.

18. Kottgen, A., Pattaro, C., Boger, C. A., Fuchsberger, C., Olden, M., Glazer, N. L., Parsa, A., Gao, X., Yang, Q., Smith, A. V., O’Connell, J. R., Li, M., Schmidt, H., Tanaka, T., Isaacs, A., Ketkar, S., Hwang, S. J., Johnson, A. D., Dehghan, A., Teumer, A., Pare, G., Atkinson, E. J., Zeller, T., Lohman, K., Cornelis, M. C., Probst-Hensch, N. M., Kronenberg, F., Tonjes, A., Hayward, C., Aspelund, T., Eiriksdottir, G., Launer, L. J., Harris, T. B., Rampersaud, E., Mitchell, B. D., Arking, D. E., Boerwinkle, E., Struchalin, M., Cavalieri, M., Singleton, A., Giallauria, F., Metter, J., de Boer, I. H., Haritunians, T., Lumley, T., Siscovick, D., Psaty, B. M., Zillikens, M. C., Oostra, B. A., Feitosa, M., Province, M., de Andrade, M., Turner, S. T., Schillert, A., Ziegler, A., Wild, P. S., Schnabel, R. B., Wilde, S., Munzel, T. F., Leak, T. S., Illig, T., Klopp, N., Meisinger, C., Wichmann, H. E., Koenig, W., Zgaga, L., Zemunik, T., Kolcic, I., Minelli, C., Hu, F. B., Johansson, A., Igl, W., Zaboli, G., Wild, S. H., Wright, A. F., Campbell, H., Ellinghaus, D., Schreiber, S., Aulchenko, Y. S., Felix, J. F., Rivadeneira, F., Uitterlinden, A. G., Hofman, A., Imboden, M., Nitsch, D., Brandstatter, A., Kollerits, B., Kedenko, L., Magi, R., Stumvoll, M., Kovacs, P., Boban, M., Campbell, S., Endlich, K., Volzke, H., Kroemer, H. K., Nauck, M., Volker, U., Polasek, O., Vitart, V., et al. (2010). New loci associated with kidney function and chronic kidney disease. Nat Genet, 42(5):376–84.

19. Lango Allen, H., Estrada, K., Lettre, G., Berndt, S. I., Weedon, M. N., Rivadeneira, F., Willer, C. J., Jackson, A. U., Vedantam, S., Raychaudhuri, S., Ferreira, T., Wood, A. R., Weyant, R. J., Segre, A. V., Speliotes, E. K., Wheeler, E., Soranzo, N., Park, J. H., Yang, J., Gudbjartsson, D., Heard-Costa, N. L., Randall, J. C., Qi, L., Vernon Smith, A., Magi, R., Pastinen, T., Liang, L., Heid, I. M., Luan, J., Thorleifsson, G., Winkler, T. W., Goddard, M. E., Sin Lo, K., Palmer, C., Workalemahu, T., Aulchenko, Y. S., Johansson, A., Zillikens, M. C., Feitosa, M. F., Esko, T., Johnson, T., Ketkar, S., Kraft, P., Mangino, M., Prokopenko, I., Absher, D., Albrecht, E., Ernst, F., Glazer, N. L., Hayward, C., Hottenga, J. J., Jacobs, K. B., Knowles, J. W., Kutalik, Z., Monda, K. L., Polasek, O., Preuss, M., Rayner, N. W., Robertson, N. R., Steinthorsdottir, V., Tyrer, J. P., Voight, B. F., Wiklund, F., Xu, J., Zhao, J. H., Nyholt, D. R., Pellikka, N., Perola, M., Perry, J. R., Surakka, I., Tammesoo, M. L., Altmaier, E. L., Amin, N., Aspelund, T., Bhangale, T., Boucher, G., Chasman, D. I., Chen, C., Coin, L., Cooper, M. N., Dixon, A. L., Gibson, Q., Grundberg, E., Hao, K., Juhani Junttila, M., Kaplan, L. M., Kettunen, J., Konig, I. R., Kwan, T., Lawrence, R. W., Levinson, D. F., Lorentzon, M., McKnight, B., Morris, A. P., Muller, M., Suh Ngwa, J., Purcell, S., Rafelt, S., Salem, R. M., Salvi, E., et al. (2010). Hundreds of variants clustered in genomic loci and biological pathways affect human height. Nature, 467(7317):832–8.

20. Lewin, A., Saadi, H., Peters, J. E., Moreno-Moral, A., Lee, J. C., Smith, K. G., Petretto, E., Bottolo, L., and Richardson, S. (2016). Mt-hess: an efficient bayesian approach for simultaneous association detection in omics datasets, with application to eqtl mapping in multiple tissues. Bioinformatics, 32(4):523–32.

21. Li, Y. and Kellis, M. (2016). Joint bayesian inference of risk variants and tissue-specific epigenomic enrichments across multiple complex human diseases. Nucleic Acids Res, 44(18):e144.

22. Li, Y. I., van de Geijn, B., Raj, A., Knowles, D. A., Petti, A. A., Golan, D., Gilad, Y., and Pritchard, J. K. (2016). Rna splicing is a primary link between genetic variation and disease. Science, 352(6285):600–4.

23. Locke, A. E., Kahali, B., Berndt, S. I., Justice, A. E., Pers, T. H., Day, F. R., Powell, C., Vedantam, S., Buchkovich, M. L., Yang, J., Croteau-Chonka, D. C., Esko, T., Fall, T., Ferreira, T., Gustafsson, S., Kutalik, Z., Luan, J., Magi, R., Randall, J. C., Winkler, T. W., Wood, A. R., Workalemahu, T., Faul, J. D., Smith, J. A., Zhao, J. H., Zhao, W., Chen, J., Fehrmann, R., Hedman, A. K., Karjalainen, J., Schmidt, E. M., Absher, D., Amin, N., Anderson, D., Beekman, M., Bolton, J. L., Bragg-Gresham, J. L., Buyske, S., Demirkan, A., Deng, G., Ehret, G. B., Feenstra, B., Feitosa, M. F., Fischer, K., Goel, A., Gong, J., Jackson, A. U., Kanoni, S., Kleber, M. E., Kristiansson, K., Lim, U., Lotay, V., Mangino, M., Leach, I. M., Medina-Gomez, C., Medland, S. E., Nalls, M. A., Palmer, C. D., Pasko, D., Pechlivanis, S., Peters, M. J., Prokopenko, I., Shungin, D., Stancakova, A., Strawbridge, R. J., Sung, Y. J., Tanaka, T., Teumer, A., Trompet, S., van der Laan, S. W., van Setten, J., Van Vliet-Ostaptchouk, J. V., Wang, Z., Yengo, L., Zhang, W., Isaacs, A., Albrecht, E., Arnlov, J., Arscott, G. M., Attwood, A. P., Bandinelli, S., Barrett, A., Bas, I. N., Bellis, C., Bennett, A. J., Berne, C., Blagieva, R., Bluher, M., Bohringer, S., Bonnycastle, L. L., Bottcher, Y., Boyd, H. A., Bruinenberg, M., Caspersen, I. H., Chen, Y. I., Clarke, R., Daw, E. W., de Craen, A. J. M., Delgado, G., Dimitriou, M., et al. (2015). Genetic studies of body mass index yield new insights for obesity biology. Nature, 518(7538):197–206.

24. MacArthur, J., Bowler, E., Cerezo, M., Gil, L., Hall, P., Hastings, E., Junkins, H., McMahon, A., Milano, A., Morales, J., Pendlington, Z. M., Welter, D., Burdett, T., Hindorff, L., Flicek, P., Cunningham, F., and Parkinson, H. (2017). The new nhgri-ebi catalog of published genome-wide association studies (gwas catalog). Nucleic Acids Res, 45(D1):D896–D901.

25. Pai, A. A., Pritchard, J. K., and Gilad, Y. (2015). The genetic and mechanistic basis for variation in gene regulation. PLoS Genet, 11(1):e1004857.

26. Pickrell, J. K., Berisa, T., Liu, J. Z., Segurel, L., Tung, J. Y., and Hinds, D. A. (2016). Detection and interpretation of shared genetic influences on 42 human traits. Nat Genet, 48(7):709–17.

27. Porter, H. F. and O’Reilly, P. F. (2017). Multivariate simulation framework reveals performance of multi-trait gwas methods. Sci Rep, 7:38837.

28. Price, A. L., Spencer, C. C., and Donnelly, P. (2015). Progress and promise in understanding the genetic basis of common diseases. Proc Biol Sci, 282(1821):20151684.

29. Sherry, S. T., Ward, M. H., Kholodov, M., Baker, J., Phan, L., Smigielski, E. M., and Sirotkin, K. (2001). dbsnp: the ncbi database of genetic variation. Nucleic Acids Res, 29(1):308–11.

30. Shi, H., Mancuso, N., Spendlove, S., and Pasaniuc, B. (2017). Local genetic correlation gives insights into the shared genetic architecture of complex traits. Am J Hum Genet, 101(5):737–751.

31. Shriner, D. (2012). Moving toward system genetics through multiple trait analysis in genome-wide association studies. Front Genet, 3:1.

32. Shungin, D., Winkler, T. W., Croteau-Chonka, D. C., Ferreira, T., Locke, A. E., Magi, R., Strawbridge, R. J., Pers, T. H., Fischer, K., Justice, A. E., Workalemahu, T., Wu, J. M. W., Buchkovich, M. L., Heard-Costa, N. L., Roman, T. S., Drong, A. W., Song, C., Gustafsson, S., Day, F. R., Esko, T., Fall, T., Kutalik, Z., Luan, J., Randall, J. C., Scherag, A., Vedantam, S., Wood, A. R., Chen, J., Fehrmann, R., Karjalainen, J., Kahali, B., Liu, C. T., Schmidt, E. M., Absher, D., Amin, N., Anderson, D., Beekman, M., Bragg-Gresham, J. L., Buyske, S., Demirkan, A., Ehret, G. B., Feitosa, M. F., Goel, A., Jackson, A. U., Johnson, T., Kleber, M. E., Kristiansson, K., Mangino, M., Leach, I. M., Medina-Gomez, C., Palmer, C. D., Pasko, D., Pechlivanis, S., Peters, M. J., Prokopenko, I., Stancakova, A., Sung, Y. J., Tanaka, T., Teumer, A., Van Vliet-Ostaptchouk, J. V., Yengo, L., Zhang, W., Albrecht, E., Arnlov, J., Arscott, G. M., Bandinelli, S., Barrett, A., Bellis, C., Bennett, A. J., Berne, C., Bluher, M., Bohringer, S., Bonnet, F., Bottcher, Y., Bruinenberg, M., Carba, D. B., Caspersen, I. H., Clarke, R., Daw, E. W., Deelen, J., Deelman, E., Delgado, G., Doney, A. S., Eklund, N., Erdos, M. R., Estrada, K., Eury, E., Friedrich, N., Garcia, M. E., Giedraitis, V., Gigante, B., Go, A. S., Golay, A., Grallert, H., Grammer, T. B., Grassler, J., Grewal, J., Groves, C. J., Haller, T., Hallmans, G., et al. (2015). New genetic loci link adipose and insulin biology to body fat distribution. Nature, 518(7538):187–196.

33. Speliotes, E. K., Willer, C. J., Berndt, S. I., Monda, K. L., Thorleifsson, G., Jackson, A. U., Allen, H. L., Lindgren, C. M., Luan, J., Magi, R., Randall, J. C., Vedantam, S., Winkler, T. W., Qi, L., Workalemahu, T., Heid, I. M., Steinthorsdottir, V., Stringham, H. M., Weedon, M. N., Wheeler, E., Wood, A. R., Ferreira, T., Weyant, R. J., Segre, A. V., Estrada, K., Liang, L., Nemesh, J., Park, J. H., Gustafsson, S., Kilpelainen, T. O., Yang, J., Bouatia-Naji, N., Esko, T., Feitosa, M. F., Kutalik, Z., Mangino, M., Raychaudhuri, S., Scherag, A., Smith, A. V., Welch, R., Zhao, J. H., Aben, K. K., Absher, D. M., Amin, N., Dixon, A. L., Fisher, E., Glazer, N. L., Goddard, M. E., Heard-Costa, N. L., Hoesel, V., Hottenga, J. J., Johansson, A., Johnson, T., Ketkar, S., Lamina, C., Li, S., Moffatt, M. F., Myers, R. H., Narisu, N., Perry, J. R., Peters, M. J., Preuss, M., Ripatti, S., Rivadeneira, F., Sandholt, C., Scott, L. J., Timpson, N. J., Tyrer, J. P., van Wingerden, S., Watanabe, R. M., White, C. C., Wiklund, F., Barlassina, C., Chasman, D. I., Cooper, M. N., Jansson, J. O., Lawrence, R. W., Pellikka, N., Prokopenko, I., Shi, J., Thiering, E., Alavere, H., Alibrandi, M. T., Almgren, P., Arnold, A. M., Aspelund, T., Atwood, L. D., Balkau, B., Balmforth, A. J., Bennett, A. J., Ben-Shlomo, Y., Bergman, R. N., Bergmann, S., Biebermann, H., Blakemore, A. I., Boes, T., Bonnycastle, L. L., Bornstein, S. R., Brown, M. J., Buchanan, T. A., et al. (2010). Association analyses of 249,796 individuals reveal 18 new loci associated with body mass index. Nat Genet, 42(11):937–48.

34. Stephens, M. (2013). A unified framework for association analysis with multiple related phenotypes. PLoS One, 8(7):e65245.

35. Stricker, S. H., Koferle, A., and Beck, S. (2017). From profiles to function in epigenomics. Nat Rev Genet, 18(1):51–66.

36. Sudlow, C., Gallacher, J., Allen, N., Beral, V., Burton, P., Danesh, J., Downey, P., Elliott, P., Green, J., Landray, M., Liu, B., Matthews, P., Ong, G., Pell, J., Silman, A., Young, A., Sprosen, T., Peakman, T., and Collins, R. (2015). Uk biobank: an open access resource for identifying the causes of a wide range of complex diseases of middle and old age. PLoS Med, 12(3):e1001779.

37. Teslovich, T. M., Musunuru, K., Smith, A. V., Edmondson, A. C., Stylianou, I. M., Koseki, M., Pirruccello, J. P., Ripatti, S., Chasman, D. I., Willer, C. J., Johansen, C. T., Fouchier, S. W., Isaacs, A., Peloso, G. M., Barbalic, M., Ricketts, S. L., Bis, J. C., Aulchenko, Y. S., Thorleifsson, G., Feitosa, M. F., Chambers, J., Orho-Melander, M., Melander, O., Johnson, T., Li, X., Guo, X., Li, M., Shin Cho, Y., Jin Go, M., Jin Kim, Y., Lee, J. Y., Park, T., Kim, K., Sim, X., Twee-Hee Ong, R., Croteau-Chonka, D. C., Lange, L. A., Smith, J. D., Song, K., Hua Zhao, J., Yuan, X., Luan, J., Lamina, C., Ziegler, A., Zhang, W., Zee, R. Y., Wright, A. F., Witteman, J. C., Wilson, J. F., Willemsen, G., Wichmann, H. E., Whitfield, J. B., Waterworth, D. M., Wareham, N. J., Waeber, G., Vollenweider, P., Voight, B. F., Vitart, V., Uitterlinden, A. G., Uda, M., Tuomilehto, J., Thompson, J. R., Tanaka, T., Surakka, I., Stringham, H. M., Spector, T. D., Soranzo, N., Smit, J. H., Sinisalo, J., Silander, K., Sijbrands, E. J., Scuteri, A., Scott, J., Schlessinger, D., Sanna, S., Salomaa, V., Saharinen, J., Sabatti, C., Ruokonen, A., Rudan, I., Rose, L. M., Roberts, R., Rieder, M., Psaty, B. M., Pramstaller, P. P., Pichler, I., Perola, M., Penninx, B. W., Pedersen, N. L., Pattaro, C., Parker, A. N., Pare, G., Oostra, B. A., O’Donnell, C. J., Nieminen, M. S., Nickerson, D. A., Montgomery, G. W., Meitinger, T., McPherson, R., McCarthy, M. I., et al. (2010). Biological, clinical and population relevance of 95 loci for blood lipids. Nature, 466(7307):707–13.

38. Urbut, S. M., Wang, G., and Stephens, M. (2017). Flexible statistical methods for estimating and testing effects in genomic studies with multiple conditions. bioRxiv, 096552.

39. van der Harst, P., Zhang, W., Mateo Leach, I., Rendon, A., Verweij, N., Sehmi, J., Paul, D. S., Elling, U., Allayee, H., Li, X., Radhakrishnan, A., Tan, S. T., Voss, K., Weichenberger, C. X., Albers, C. A., Al-Hussani, A., Asselbergs, F. W., Ciullo, M., Danjou, F., Dina, C., Esko, T., Evans, D. M., Franke, L., Gogele, M., Hartiala, J., Hersch, M., Holm, H., Hottenga, J. J., Kanoni, S., Kleber, M. E., Lagou, V., Langenberg, C., Lopez, L. M., Lyytikainen, L. P., Melander, O., Murgia, F., Nolte, I. M., O’Reilly, P. F., Padmanabhan, S., Parsa, A., Pirastu, N., Porcu, E., Portas, L., Prokopenko, I., Ried, J. S., Shin, S. Y., Tang, C. S., Teumer, A., Traglia, M., Ulivi, S., Westra, H. J., Yang, J., Zhao, J. H., Anni, F., Abdellaoui, A., Attwood, A., Balkau, B., Bandinelli, S., Bastardot, F., Benyamin, B., Boehm, B. O., Cookson, W. O., Das, D., de Bakker, P. I., de Boer, R. A., de Geus, E. J., de Moor, M. H., Dimitriou, M., Domingues, F. S., Doring, A., Engstrom, G., Eyjolfsson, G. I., Ferrucci, L., Fischer, K., Galanello, R., Garner, S. F., Genser, B., Gibson, Q. D., Girotto, G., Gudbjartsson, D. F., Harris, S. E., Hartikainen, A. L., Hastie, C. E., Hedblad, B., Illig, T., Jolley, J., Kahonen, M., Kema, I. P., Kemp, J. P., Liang, L., Lloyd-Jones, H., Loos, R. J., Meacham, S., Medland, S. E., Meisinger, C., Memari, Y., Mihailov, E., Miller, K., Moffatt, M. F., Nauck, M., et al. (2012). Seventy-five genetic loci influencing the human red blood cell. Nature, 492(7429):369–75.

40. Visscher, P. M., Wray, N. R., Zhang, Q., Sklar, P., McCarthy, M. I., Brown, M. A., and Yang, J. (2017). 10 years of gwas discovery: Biology, function, and translation. Am J Hum Genet, 101(1):5–22.

41. Wain, L. V., Verwoert, G. C., O’Reilly, P. F., Shi, G., Johnson, T., Johnson, A. D., Bochud, M., Rice, K. M., Henneman, P., Smith, A. V., Ehret, G. B., Amin, N., Larson, M. G., Mooser, V., Hadley, D., Dorr, M., Bis, J. C., Aspelund, T., Esko, T., Janssens, A. C., Zhao, J. H., Heath, S., Laan, M., Fu, J., Pistis, G., Luan, J., Arora, P., Lucas, G., Pirastu, N., Pichler, I., Jackson, A. U., Webster, R. J., Zhang, F., Peden, J. F., Schmidt, H., Tanaka, T., Campbell, H., Igl, W., Milaneschi, Y., Hottenga, J. J., Vitart, V., Chasman, D. I., Trompet, S., Bragg-Gresham, J. L., Alizadeh, B. Z., Chambers, J. C., Guo, X., Lehtimaki, T., Kuhnel, B., Lopez, L. M., Polasek, O., Boban, M., Nelson, C. P., Morrison, A. C., Pihur, V., Ganesh, S. K., Hofman, A., Kundu, S., Mattace-Raso, F. U., Rivadeneira, F., Sijbrands, E. J., Uitterlinden, A. G., Hwang, S. J., Vasan, R. S., Wang, T. J., Bergmann, S., Vollenweider, P., Waeber, G., Laitinen, J., Pouta, A., Zitting, P., McArdle, W. L., Kroemer, H. K., Volker, U., Volzke, H., Glazer, N. L., Taylor, K. D., Harris, T. B., Alavere, H., Haller, T., Keis, A., Tammesoo, M. L., Aulchenko, Y., Barroso, I., Khaw, K. T., Galan, P., Hercberg, S., Lathrop, M., Eyheramendy, S., Org, E., Sober, S., Lu, X., Nolte, I. M., Penninx, B. W., Corre, T., Masciullo, C., Sala, C., Groop, L., Voight, B. F., Melander, O., et al. (2011). Genome-wide association study identifies six new loci influencing pulse pressure and mean arterial pressure. Nat Genet, 43(10):1005–11.

42. Wen, X., Pique-Regi, R., and Luca, F. (2017). Integrating molecular qtl data into genome-wide genetic association analysis: Probabilistic assessment of enrichment and colocalization. PLoS Genet, 13(3):e1006646.

43. Willer, C. J., Schmidt, E. M., Sengupta, S., Peloso, G. M., Gustafsson, S., Kanoni, S., Ganna, A., Chen, J., Buchkovich, M. L., Mora, S., Beckmann, J. S., Bragg-Gresham, J. L., Chang, H. Y., Demirkan, A., Den Hertog, H. M., Do, R., Donnelly, L. A., Ehret, G. B., Esko, T., Feitosa, M. F., Ferreira, T., Fischer, K., Fontanillas, P., Fraser, R. M., Freitag, D. F., Gurdasani, D., Heikkila, K., Hypponen, E., Isaacs, A., Jackson, A. U., Johansson, A., Johnson, T., Kaakinen, M., Kettunen, J., Kleber, M. E., Li, X., Luan, J., Lyytikainen, L. P., Magnusson, P. K. E., Mangino, M., Mihailov, E., Montasser, M. E., Muller-Nurasyid, M., Nolte, I. M., O’Connell, J. R., Palmer, C. D., Perola, M., Petersen, A. K., Sanna, S., Saxena, R., Service, S. K., Shah, S., Shungin, D., Sidore, C., Song, C., Strawbridge, R. J., Surakka, I., Tanaka, T., Teslovich, T. M., Thorleifsson, G., Van den Herik, E. G., Voight, B. F., Volcik, K. A., Waite, L. L., Wong, A., Wu, Y., Zhang, W., Absher, D., Asiki, G., Barroso, I., Been, L. F., Bolton, J. L., Bonnycastle, L. L., Brambilla, P., Burnett, M. S., Cesana, G., Dimitriou, M., Doney, A. S. F., Doring, A., Elliott, P., Epstein, S. E., Ingi Eyjolfsson, G., Gigante, B., Goodarzi, M. O., Grallert, H., Gravito, M. L., Groves, C. J., Hallmans, G., Hartikainen, A. L., Hayward, C., Hernandez, D., Hicks, A. A., Holm, H., Hung, Y. J., Illig, T., Jones, M. R., Kaleebu, P., Kastelein, J. J. P., Khaw, K. T., Kim, E., et al. (2013). Discovery and refinement of loci associated with lipid levels. Nat Genet, 45(11):1274–1283.

44. Wood, A. R., Esko, T., Yang, J., Vedantam, S., Pers, T. H., Gustafsson, S., Chu, A. Y., Estrada, K., Luan, J., Kutalik, Z., Amin, N., Buchkovich, M. L., Croteau-Chonka, D. C., Day, F. R., Duan, Y., Fall, T., Fehrmann, R., Ferreira, T., Jackson, A. U., Karjalainen, J., Lo, K. S., Locke, A. E., Magi, R., Mihailov, E., Porcu, E., Randall, J. C., Scherag, A., Vinkhuyzen, A. A., Westra, H. J., Winkler, T. W., Workalemahu, T., Zhao, J. H., Absher, D., Albrecht, E., Anderson, D., Baron, J., Beekman, M., Demirkan, A., Ehret, G. B., Feenstra, B., Feitosa, M. F., Fischer, K., Fraser, R. M., Goel, A., Gong, J., Justice, A. E., Kanoni, S., Kleber, M. E., Kristiansson, K., Lim, U., Lotay, V., Lui, J. C., Mangino, M., Mateo Leach, I., Medina-Gomez, C., Nalls, M. A., Nyholt, D. R., Palmer, C. D., Pasko, D., Pechlivanis, S., Prokopenko, I., Ried, J. S., Ripke, S., Shungin, D., Stancakova, A., Strawbridge, R. J., Sung, Y. J., Tanaka, T., Teumer, A., Trompet, S., van der Laan, S. W., van Setten, J., Van VlietOstaptchouk, J. V., Wang, Z., Yengo, L., Zhang, W., Afzal, U., Arnlov, J., Arscott, G. M., Bandinelli, S., Barrett, A., Bellis, C., Bennett, A. J., Berne, C., Bluher, M., Bolton, J. L., Bottcher, Y., Boyd, H. A., Bruinenberg, M., Buckley, B. M., Buyske, S., Caspersen, I. H., Chines, P. S., Clarke, R., Claudi-Boehm, S., Cooper, M., Daw, E. W., De Jong, P. A., Deelen, J., Delgado, G., et al. (2014). Defining the role of common variation in the genomic and biological architecture of adult human height. Nat Genet, 46(11):1173–86.

45. Yang, Q. and Wang, Y. (2012). Methods for analyzing multivariate phenotypes in genetic association studies. J Probab Stat, 2012:652569.

46. Zheng, H. F., Forgetta, V., Hsu, Y. H., Estrada, K., Rosello-Diez, A., Leo, P. J., Dahia, C. L., Park-Min, K. H., Tobias, J. H., Kooperberg, C., Kleinman, A., Styrkarsdottir, U., Liu, C. T., Uggla, C., Evans, D. S., Nielson, C. M., Walter, K., Pettersson-Kymmer, U., McCarthy, S., Eriksson, J., Kwan, T., Jhamai, M., Trajanoska, K., Memari, Y., Min, J., Huang, J., Danecek, P., Wilmot, B., Li, R., Chou, W. C., Mokry, L. E., Moayyeri, A., Claussnitzer, M., Cheng, C. H., Cheung, W., Medina-Gomez, C., Ge, B., Chen, S. H., Choi, K., Oei, L., Fraser, J., Kraaij, R., Hibbs, M. A., Gregson, C. L., Paquette, D., Hofman, A., Wibom, C., Tranah, G. J., Marshall, M., Gardiner, B. B., Cremin, K., Auer, P., Hsu, L., Ring, S., Tung, J. Y., Thorleifsson, G., Enneman, A. W., van Schoor, N. M., de Groot, L. C., van der Velde, N., Melin, B., Kemp, J. P., Christiansen, C., Sayers, A., Zhou, Y., Calderari, S., van Rooij, J., Carlson, C., Peters, U., Berlivet, S., Dostie, J., Uitterlinden, A. G., Williams, S. R., Farber, C., Grinberg, D., LaCroix, A. Z., Haessler, J., Chasman, D. I., Giulianini, F., Rose, L. M., Ridker, P. M., Eisman, J. A., Nguyen, T. V., Center, J. R., Nogues, X., Garcia-Giralt, N., Launer, L. L., Gudnason, V., Mellstrom, D., Vandenput, L., Amin, N., van Duijn, C. M., Karlsson, M. K., Ljunggren, O., Svensson, O., Hallmans, G., Rousseau, F., Giroux, S., Bussiere, J., Arp, P. P., et al. (2015). Whole-genome sequencing identifies en1 as a determinant of bone density and fracture. Nature, 526(7571):112–7.

47. Zhu, W. and Zhang, H. (2009). Why do we test multiple traits in genetic association studies? J Korean Stat Soc, 38(1):1–10.

48. Zhu, Z., Zhang, F., Hu, H., Bakshi, A., Robinson, M. R., Powell, J. E., Mont-gomery, G. W., Goddard, M. E., Wray, N. R., Visscher, P. M., and Yang, J. (2016). Integration of summary data from gwas and eqtl studies predicts complex trait gene targets. Nat Genet, 48(5):481–7.

