## Supplementary Figures & Tables for "Bayesian multivariate reanalysis of large genetic studies identifies many new associations"

August 1, 2019

### 1 Supplementary Figures

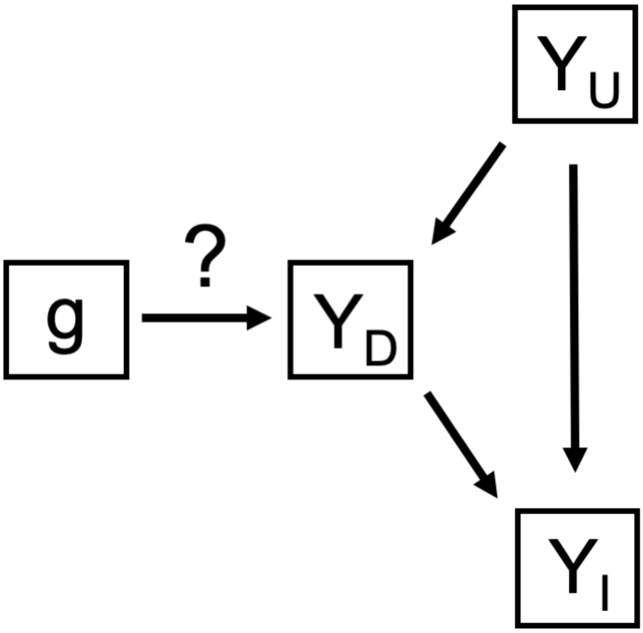


Supplementary Figure 1: **Graphical Model of Multivariate Categories**. Shown here is a Directed Acyclic Graphical (DAG) model of our multivariate categories in the context of our vector of phenotypes $\mathbf{Y}$ (e.g. $\mathbf{Y} = \{\mathbf{Y_U}, \mathbf{Y_D}, \mathbf{Y_I}\}$) and their connections with the variant of interest $\mathbf{g}$. The relationships described in-text can be seen here. $\mathbf{Y_U}$, our unassociated phenotypes, have no connection with $\mathbf{g}$. $\mathbf{Y_D}$, our directly associated phenotypes, have a direct connection with $\mathbf{g}$. And $\mathbf{Y_I}$, our indirectly associated phenotypes, have a connection with $\mathbf{g}$ only by going through $\mathbf{Y_D}$ first. Note that if $\mathbf{Y_D}$ were not observed, $\mathbf{Y_I}$ would appear as a direct connection.

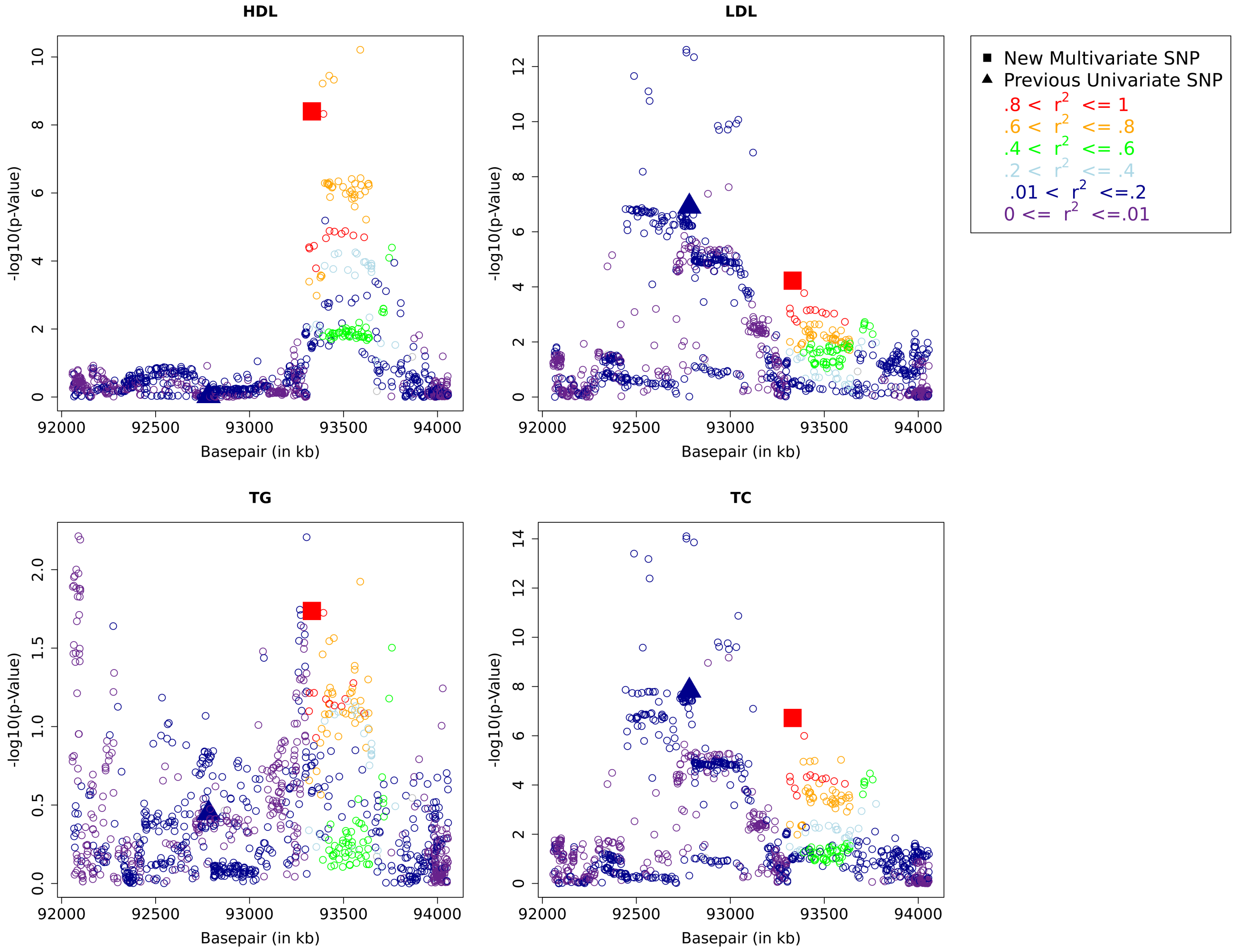


Supplementary Figure 2: **Refining Association Signals – GlobalLipids2013 rs7515577 & rs12038699**. Shown are the -$\log_{10}$ univariate *p*-values from the GlobalLipids2013 analysis for both the previous univariate association rs7515577 ("Previous Univariate SNP") and the new multivariate association rs12038699 ("New Multivariate SNP") across all four phenotypes analyzed. rs7515577 is represented as a triangle and rs12038699 is represented as a square. Also shown are the -$\log_{10}$ univariate *p*-values of SNPs within 1Mb of the midpoint between rs7515577 and rs12038699. Color-coding of the SNPs represent the degree of linkage disequilibrium between variants and the new association rs12038699 based on the GBR cohort of 1000Genomes (Genomes Project et al., 2015); for color coding details, see legend.

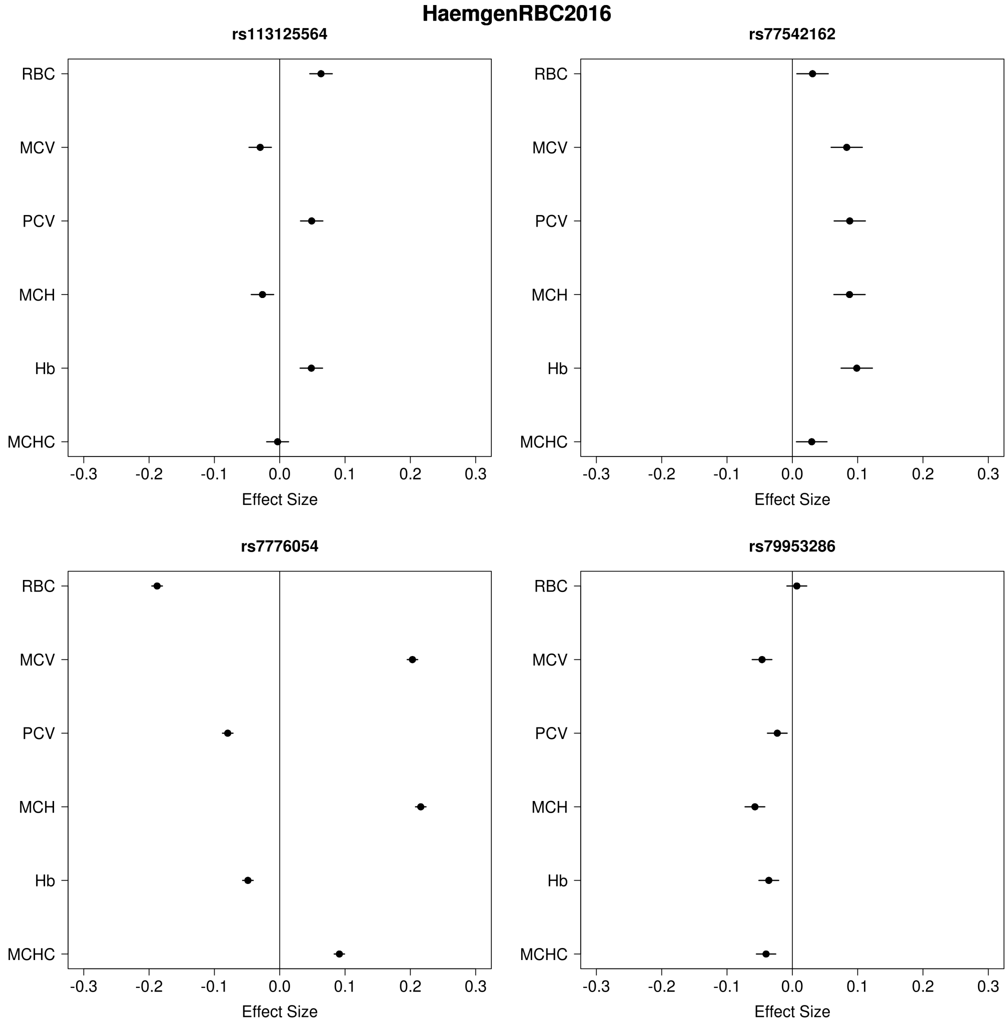


Supplementary Figure 3: **Effect Size Heterogeneity Among SNPs With Identical Multivariate Model Assignments**. Shown are the phenotype effect sizes (points), and ±2 standard errors (bars), for four significantly associated SNPs from HaemgenRBC2016. All four SNPs were classified as being "associated" with all six phenotypes (i.e. marginal posterior probability of association >= 95% for each phenotype). However, they clearly show different patterns of effect sizes. Therefore focusing simply on binary calls of "associated" vs "unassociated" can hide different patterns of multivariate association.

### 2 Supplementary Tables

Supplementary Table 1: **Summary of Associations in Each Dataset**

| Dataset | Release | New Multivariate SNPs[a] | Univariate $p$-Value Threshold[b] | New SNPs Below Univariate Threshold[c] | New SNPs Near Previous Association[d] |
|---|---|---|---|---|---|
| GlobalLipids | 2010 | 19 | $5 \times 10^{-8}$ | 0 | 0 |
| | 2013 | 65 | $5 \times 10^{-8}$ | 18 | 11 |
| GIANT | 2010 | 60 | $5 \times 10^{-8}$ | 6 | 5 |
| | 2014/5 | 162 | $5 \times 10^{-8}$ | 41 | 21 |
| HaemgenRBC | 2012 | 16 | $1 \times 10^{-8}$ | 4 | 3 |
| | 2016 | 60 | $8.31 \times 10^{-9}$ | 0 | 0 |
| ICBP | 2011 | 22 | $5 \times 10^{-8}$ | 2 | 1 |
| MAGIC | 2010 | 1 | $5 \times 10^{-8}$ | 0 | 0 |
| GEFOS | 2015 | 13 | $1.2 \times 10^{-8}$ | 3 | 1 |
| GIS | 2014 | 5 | $5 \times 10^{-8}$ | 5 | 0 |
| SSGAC | 2016 | 1 | $5 \times 10^{-8}$ | 0 | 0 |
| CKDGen | 2010/1 | 6 | $5 \times 10^{-8}$ | 0 | 0 |
| ENIGMA2 | 2015 | 3 | $7.1 \times 10^{-9}$ | 0 | 0 |

[a] Number of new multivariate associations discovered by our analysis. Note that we required a multivariate association to be at least 500kb from a previous reported association to be considered “new”.

[b] Univariate GWAS significance $p$-value threshold used by the original study publication.

[c] These are new multivariate SNPs that were not reported by the original study despite having a univariate association (in the public summary data) that was genome-wide significant by the original study’s univariate significance threshold.

[d] A “previous association” means an association reported by the original GWAS; “near” means within 1Mb (but these are all more than 500kb away from a previous association since our classification of new multivariate SNPs requires this).

Supplementary Tables 2a-m: **Lists of New `bmass` Multivariate Associations, per Dataset**. Attached Excel sheets list new `bmass` associations for each dataset analyzed.

Supplementary Tables 3a-m: **Lists of Retrieved Univariate Associations From Original Publications, per Dataset**. Attached Excel sheets list the rsID#'s of the univariate significant SNPs that were retrieved from the original publication(s) associated with each dataset (see Online Methods for details).

Supplementary Tables 4a-m: **Results for Previous Univariate Associations, per Dataset**. Attached Excel sheets give `bmass` results for previous univariate associations, per dataset. Note that these results may not include all SNPs from Tables 3a-m, because some SNPs were dropped during QC and other SNPs were dropped because they did not reach univariate significance in the publicly available summary data (see Online Methods for details).

| **Dataset** | **Releases ($1^{st}$, $2^{nd}$)** | **Univariate *p*-Value Threshold ($1^{st}$, $2^{nd}$)** | **New Multivariate SNPs in $1^{st}$** | **Lower Univariate *p*-Value in $2^{nd}$** | **Below $2^{nd}$ Univariate Threshold** |
|---|---|---|---|---|---|
| GlobalLipids | 2010, 2013 | $5 \times 10^{-8}$ ,$5 \times 10^{-8}$ | 19 | 14 | 11 |
| GIANT | 2010, 2014/5 | $5 \times 10^{-8}$, $5 \times 10^{-8}$ | 60 | 59 | 51 |
| HaemgenRBC | 2012, 2016 | $1 \times 10^{-8}$, $8.31 \times 10^{-9}$ | 16 | 11 | 6 |

Supplementary Table 5: **Replication of New Multivariate Associations**. Shown are example metrics of how well our new multivariate associations replicate in datasets that allow such an evaluation. Specifically, for three of the studies used (GlobalLipids, GIANT, and HaemgenRBC), there are multiple dataset releases. To examine how well our new multivariate `bmass` associations replicate, we compared the results from the first releases ("$1^{st}$") with the univariate GWAS associations of the second releases ("$2^{nd}$"). In essence, each of these approaches aim to increase power – one by using a multivariate approach (`bmass`) and the other by increasing sample size (the $2^{nd}$ releases) – thus allowing us to compare the results against one another. Univariate *p*-Value Threshold: univariate GWAS significance *p*-value thresholds used by the original publication(s) for both the earlier ($1^{st}$) and later ($2^{nd}$) releases. New Multivariate SNPs in $1^{st}$: number of new multivariate associations from the earlier release. Lower Univariate *p*-Value in $2^{nd}$: number of new multivariate associations from the earlier release that also have lower univariate *p*-values in the later release. Below $2^{nd}$ Univariate Threshold: number of new multivariate associations from the earlier release that also cross the later release's univariate GWAS significance threshold.

| SNP | Phenotype | Direction[a] | 2010[b] | 2013[c] |
|---|---|---|---|---|
| **rs7515577** | | | | |
| (Previous) | HDL | + | 9.81E-01 | 9.29E-01 |
| | LDL | - | 1.51E-07 | 1.21E-07 |
| | TG | - | 1.80E-01 | 3.57E-01 |
| | TC | - | 2.78E-08 | 1.47E-08 |
| **rs12038699** | | | | |
| (New) | HDL | + | 4.22E-05 | 3.98E-09 |
| | LDL | + | 1.06E-03 | 5.95E-05 |
| | TG | + | 8.51E-02 | 1.83E-02 |
| | TC | - | 7.12E-05 | 1.90E-07 |

[a] Whether the reference allele increases (+) or decreases (-) phenotype.
[b] $p$-value from GlobalLipids 2010.
[c] $p$-value from GlobalLipids 2013.

Supplementary Table 6: **$p$-Values for rs7515577 & rs12038699 in 2010 and 2013 GlobalLipds Releases** – In the 2010 release rs7515577 has a univariate $p$-value that crosses the $5 \times 10^{-8}$ threshold (TC), whereas rs12038699 does not. Since rs12038699 is near to rs7515577 it may get masked for future analyses; however in the 2013 data rs12038699 not only has a lower minimum univariate $p$-value, but also has a different multivariate $p$-value pattern as compared to rs7515577. Both these signals suggest that rs12038699 should be viewed as a separate GWAS hit for GlobalLipids2013.

**GlobalLipids2013: New+Prev SNPs**

| HDL_LDL TG_TC | n | Mean Posterior | Original Prior |
|---|---|---|---|
| 1_2_1_1 | 100 | 0.635 | 0.327 |
| 1_2_2_1 | 38 | 0.531 | 0.138 |
| 1_1_1_2 | 18 | 0.751 | 0.117 |
| 2_1_2_2 | 17 | 0.661 | 0.057 |
| 1_1_1_1 | 11 | 0.769 | 0.078 |

**GIANT2014/5: New+Prev SNPs**

| Height_BMI WHRadjBMI | n | Mean Posterior | Original Prior |
|---|---|---|---|
| 1_2_0 | 616 | 0.443 | 0.318 |
| 1_1_1 | 143 | 0.677 | 0.161 |
| 1_1_0 | 89 | 0.488 | 0.094 |
| 1_2_1 | 15 | 0.387 | 0.037 |
| 1_2_2 | 13 | 0.623 | 0.257 |

**HaemgenRBC2016: New+Prev SNPs**

| RBC_MCV_PCV MCH_Hb_MCHC | n | Mean Posterior | Original Prior |
|---|---|---|---|
| 2_1_1_2_2_2 | 179 | 0.487 | 0.17 |
| 2_1_2_2_1_1 | 162 | 0.502 | 0.203 |
| 2_1_0_2_2_1 | 105 | 0.498 | 0.117 |
| 2_1_1_2_2_1 | 51 | 0.561 | 0.155 |
| 2_0_1_2_2_2 | 33 | 0.405 | 0.038 |

Supplementary Table 7: **Top Multivariate Model Examples per SNP**. List of multivariate models that most frequently have the highest posterior probabilities per SNP. Top 5 models are shown from across both the previous univariate associations analyzed and the new multivariate associations discovered in the GlobalLipids2013, GIANT2014/5, and HaemgenRBC2016 datasets. Phenotype ordering is shown in the header, where 0, 1, and 2 refer to the multivariate categories of **U**nassociated, **D**irectly Associated, and **I**ndirectly Associated. n is the number of SNPs that show the specified model as having the largest posterior probability, with Mean Posterior displaying the average posterior probability of the given model across the n SNPs, and Original Prior showing the prior established for the given model from training on all the previous univariate associations from that dataset.

### Supplementary References

Genomes Project, C., Auton, A., Brooks, L. D., Durbin, R. M., Garrison, E. P., Kang, H. M., Korbel, J. O., Marchini, J. L., McCarthy, S., McVean, G. A., and Abecasis, G. R. (2015). A global reference for human genetic variation. *Nature*, 526(7571):68–74.
